# A Deep Learning-Based Segmentation of Cells and Analysis (DL-SCAN)

**DOI:** 10.1101/2024.05.03.592244

**Authors:** Alok Bhattarai, Jan Meyer, Laura Petersilie, Syed I Shah, Christine R. Rose, Ghanim Ullah

## Abstract

With the recent surge in the development of highly selective probes, fluorescence microscopy has become one of the most widely used approaches to study cellular properties and signaling in living cells and tissues. Traditionally, microscopy image analysis heavily relies on manufacturer-supplied software, which often demands extensive training and lacks automation capabilities for handling diverse datasets. A critical challenge arises, if fluorophores employed exhibit low brightness and low Signal-to-Noise ratio (SNR). As a consequence, manual intervention may become a necessity, introducing variability in the analysis outcomes even for identical samples when analyzed by different users. This leads to the incorporation of blinded analysis which ensures that the outcome is free from user bias to a certain extent but is extremely time-consuming. To overcome these issues, we have developed a tool called DL-SCAN that automatically segments and analyzes fluorophore-stained regions of interest such as cell bodies in fluorescence microscopy images using a Deep Learning algorithm called Stardist. We demonstrate the program’s ability to automate cell identification and study cellular ion dynamics using synthetic image stacks with varying SNR. This is followed by its application to experimental Na^+^ and Ca^2+^ imaging data from neurons and astrocytes in mouse brain tissue slices exposed to transient chemical ischemia. The results from DL-SCAN are consistent, reproducible, and free from user bias, allowing efficient and rapid analysis of experimental data in an objective manner. The open-source nature of the tool also provides room for modification and extension to analyze other forms of microscopy images specific to the dynamics of different ions in other cell types.

**Statement of Significance:** Fluorescence microscopy is widely used to study the functional and morphological features of living cells. However, various factors, such as low SNR, background noise, drift in the signal, movement of the tissue, and the large size of the resulting imaging data, make the processing of fluorescence microscopy data prone to errors, user bias, and extremely time-consuming. These and other issues hinder the full utilization of these powerful experimental techniques. Our novel Deep Learning-based tool overcomes these issues by processing and analyzing fluorescence imaging data, e.g., enabling automated visualization of ion changes in living cells in brain slices. Yet the tool remains easy to use with a streamlined workflow.

## Introduction

Recent advancements in imaging technology for studying biological processes at scales ranging from single molecule to the cell network and tissue level have revolutionized biological research. In particular, fluorescence microscopy techniques are widely used to study the structure and/or function of cells, or spatiotemporal changes thereof, under a variety of conditions [1-6]. These experiments generate huge image-based datasets, which may have many different objects (e.g., molecules, organelles, and cells) that need to be analyzed. Typically, this kind of analysis relies heavily on manual or semi-automated approaches, which are labor intensive, costly, not very accurate, and often poorly reproducible. Thus, fully automated methods that can overcome these issues are key to utilizing the full potential of these powerful experimental techniques.

An additional problem is the dynamic nature of the biological processes under study, which makes the analysis of imaging data even more cumbersome. For example, in many organs such as the brain, the concentration of the molecules of interest, e.g. of ions such as Ca^2+^ and Na^+^, undergoes transient changes both in time and space [7-9], while tissue movements additionally shift the exact location of the cells or cellular compartments in the field of view. In such a scenario, the user is often required to draw or adjust the regions of interest (ROIs) in which fluorescence signals are to be analyzed frame by frame. In extreme conditions, such as ischemia and seizures, brain cells swell significantly, causing the tissue to move both in the x-y and the z-axis [10, 11]. In those experiments, proper analysis of fluorescence signals is even more time-consuming. This makes the use of accurate automated methods even more desirable.

Chemical fluorescent indicators or genetically-encoded sensors are routinely used to study the spatial and/or temporal dynamics of various ions including Ca^2+^ and Na^+^. The fluorescence emission of these fluorophores can be examined using different microscopy techniques after being introduced into the sample [12]. Microscope vendors have developed several tools to serve this purpose (e.g. NIS Elements 6.0 from Nikon Europe B.V., Amstelven, the Netherlands, or Cell Sense FluoView 3.1.1 from Evident Europe GmbH, Hamburg, Germany). These tools rely mainly on manual selection of ROIs and additional programs for data evaluation are often required to assess the images and experimental datasets in more detail. For imaging of changes in intracellular Ca^2+^, for example, several tools emerged recently (e.g. AQuA, CaSCaDe, MSparkles) which automatically select ROIs based on changes in the intensity of neighboring pixels [13-16]. Most of these tools were developed with a specific application in mind, such as dynamic Ca^2+^ imaging *in vivo*. In addition, many of them rely on the use of MATLAB and commercial toolboxes, and require extensive knowledge of MATLAB as well as prior coding experience.

To overcome the above-mentioned drawbacks, we have developed a tool, called DL-SCAN. DL-SCAN is based on Python and is available as open-source. One of the development goals of DL-SCAN was to ensure usability without extensive training or prior knowledge of Python as a programming language. This is achieved through a simplified user experience with a browser-based interface. DL-SCAN takes microscopy image time series data as an input, applies a Deep Learning algorithm to segment fluorescently labeled objects and then extracts statistics about the dynamics of ions under normal and pathological states. A range of options for pre-processing and post-processing are also provided. The automated nature of DL-SCAN is designed to mitigate bias and reduce analysis time, ensuring verifiable and reproducible results. Furthermore, we posit that the development of open-source tools, such as ours, will be beneficial in the study of ion dynamics under various physiological and pathological conditions. Finally, the open-source and user-friendly nature of our approach makes extending and customizing our program for other experimental conditions or needs straightforward.

## Materials and Methods

To develop an automated tool that pre-processes, segments, and extracts various properties of time-lapse microscopy image stacks (in TIFF format, but other formats can be easily added), we made use of Python 3.8.8 and a python library called Streamlit (version 1.21.0). This allowed the development of a clean and user-friendly graphical user interface (GUI) without HTML, CSS, and JS. Here, we outline the key characteristics of DL-SCAN.

### Algorithm and Key Features of the Application

DL-SCAN initiates within a browser as a local host upon following the steps as mentioned in the User Manual (Supplementary Information Text). With “Pre-processing and Segmentation” being the homepage of the tool, it allows the user to upload a time-lapse microscopy image stack. As soon as the file is uploaded, the program checks whether the pixel values are already in unsigned integer 8 (uint8) format. If not, it performs the conversion, and subsequently, the page displays each image frame of the original stack as selected by the user. To speed up the upload and the following processes, initial conversion to uint8 format before uploading is recommended. Depending on the quality of the images generated due to variations in the experimental setup, various pre-processing techniques might be necessary to segment, characterize, and analyze them with a high level of accuracy. This tool provides multiple pre-processing options to the user that include background correction, Gaussian blur, median blur, brightness and contrast adjustments and Contrast Limited Adaptive Histogram Equalization (CLAHE).

Background subtraction is often required to remove signals not generated by the fluorophores being analyzed, such as tissue autofluorescence. Upon selecting “Background Correction”, the user can draw a single rectangle on the first image frame and select a region apparently free of distinct cellular fluorescence. At the backend, the code then computes the average intensity of the pixels in the selected background region and subtracts it from each pixel value across all frames to achieve the desired correction. The background-corrected frames will be displayed immediately after the region selection. It should be noted that these background-corrected images are not used to segment the cells, but to extract their intensity profiles to aid robust analysis. The processed frames as a result of user-selected preprocessing options are then displayed.

To improve accuracy for robust segmentation, DL-SCAN collapses all the frames into one and displays the collapsed image. This process ensures that all ROIs are properly identified and segmented once the Deep Learning segmentation algorithm is applied. The displayed collapsed image is now ready for segmentation. However, users can also choose to use the collapsed image or the first image in the stack for segmentation. The rolling ball background subtraction (RBBS) option is also incorporated at this stage of the program (Supplementary Information Text). Once the “Segment and Generate Labels” button is clicked, DL-SCAN applies a Deep Learning algorithm to identify and segment the objects. The segmented and labeled objects are then displayed. The tool is now prepared for frame-by-frame image analysis, contingent upon the identification and segmentation of the objects.

**Stardist** is a Convolutional Neural Network (CNN) Deep Learning architecture that works remarkably well in detection and segmentation of star-convex polygon shapes [17]. In other words, objects that exhibit visible boundaries from their centers are effectively predicted using Stardist [17]. The algorithm relies on U-Net based framework specifically designed to segment nuclei and cells in microscopy images, as the model is pre-trained on diverse fluorescent microscopy images of nuclei [18]. The model integrates object probability prediction and distances to the object boundary along a predetermined set of radial directions referred to as rays. The final result is achieved by applying non-maximum suppression (NMS), which is a technique to filter out overlapping or redundant object detections, determined by a set threshold. Given our focus on the detection and segmentation of neuronal and glial cells during the course of the tool development, we leveraged Stardist, which yielded favorable results without necessitating any modifications to the original training dataset.

Although DL-SCAN detects and identifies the star-convex polygon shapes exceptionally well, some cells in the image may sometimes remain unidentified, often due to the poor image quality or characteristics. This can be mitigated by using various preprocessing options. It is recommended to utilize FIJI for a broader range of preprocessing options in case the capabilities of the in-tool options fall short of a specific requirement. However, in the instances where it is crucial, users also have the additional option to manually draw ROIs on the labeled collapsed image. These hand-drawn regions get automatically added to the list of labels for further analysis.

Upon cell detection, users can utilize the “Single-cell Analysis” tab located in the sidebar to delve deep into the behavior of detected cells at an individual cell level. To enhance accessibility and streamline referencing, the selected (either the first or the collapsed), segmented and labeled images from “Preprocessing and Segmentation” are once again presented within the “Single-cell Analysis” tab. Furthermore, downloadable, and interactive tables are displayed separately, featuring mean intensity values and counts of bright pixel values surpassing a specific threshold for each identified cell across all frames. This threshold is user-adjustable and is referred to as area threshold percentage. When segmented on the collapsed image, e.g. obtained from microscopy data for cells stained with cell-permeant dyes like Calcein or Fura-2 AM [19-21], its purpose is to account for the observed variations in cell areas across different frames which effectively captures the phenomenon of spatial spread of the signal or cell swelling/shrinking with time.

Within the interactive table, users have the option to choose a single identified cell for in-depth analysis. Upon selecting the cell, an image with the highlighted chosen cell is presented, followed by a range of adjustable options. The “Frame Rate” widget enables users to input a value that converts frame numbers into time in seconds, based on the recording frequency in the experiment. Moreover, users can choose to conduct an analysis with or without bleaching correction (Supplementary Information Text), depending on the experimental setup and methodology employed. In the absence of bleaching correction, users are presented with choices to fine-tune the moving average for trace smoothing. This is followed by the option to select “Static” or “Dynamic” analysis. The “Single Frame Value” under the “Static” option allows users to input the baseline frame number (starting frame/reference frame), peak intensity frame number (frame of maximum activity), and the signal recovery frame number (frame where the signal returns to normal). On the other hand, the “Average Frame Value” differs from the previous only in a sense that it asks users to select the number of consecutive frames to calculate their average intensity values as the baseline intensity. Here, frame refers to an individual, captured image within the stack that corresponds to a specific time point, and the choice of single or average frame value depends entirely on the nature of the experiment.

With the “Dynamic” option selected, the program asks the user to select frame numbers for calculating baseline intensity using their mean intensity values, but the peak intensity and the recovery point are computed automatically by the algorithm. Subsequently, the normalized intensity table relative to the baseline, and various traces (Mean Intensity Vs. Time, Smoothed Mean Intensity Vs. Time, and Bright Pixel Area Vs. Time) are displayed. With bleaching correction enabled, several options remain consistent as previously mentioned. Photobleaching, which refers to the gradual loss of fluorescence from a fluorophore over time due to light exposure, is a common challenge in microscopy. To account for this, additionally, user-adjustable settings are introduced to select several of the first and the last frames for mono-exponential decay fitting. The intermediate data points are interpolated, resulting in the generation of the entire dataset that represents photobleaching. This is followed by their subtraction from the original data points, which leads to the production of data corrected for photobleaching. As is the case for the absence of photobleaching correction option, the normalized intensity table in relation to the baseline, and various traces mentioned above are displayed for the latter as well.

Further analysis is carried out on the “Smoothed Mean Intensity Vs. Time” trace to obtain some key parameters of changes in fluorescence intensity detected upon a specific experimental manipulation or stimulation in the chosen ROIs. These include rise time, decay time, duration, rise rate, decay rate, and peak amplitude of the induced signals. Upon clicking the “Obtain the parameters for selected label” button, these values are presented in a downloadable table.

Within the “Multi-cell Analysis” tab located in the sidebar, the user has the ability to collectively analyze multiple cells. After selecting labels from the interactive table, traces for each selected cell are displayed. These traces can be toggled on and off by clicking the corresponding legend denoted by the respective label number. To isolate the trace of interest, a double-click action is required. Subsequently, the process employs all the options as outlined in the context of single-cell analysis. The resulting table, containing the properties of the chosen cells, is downloadable, making it available for further utilization. All these properties of the tool are comprehensively outlined in the User Manual (Supplementary Information Text).

#### Synthetic Data

To test the effectiveness and robustness of the program before applying it to the real biological data, we first generated stacks of synthetic data resembling neuronal microscopy images (150 frames each in TIFF format) using Python. Single and multiple object datasets were produced across a range of SNR values. The datasets were created using a simple approach defined by the formula:

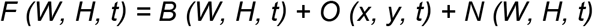

Here, *F* is the final image for the *t*^*th*^ frame with width *W* and height *H*, and is a result of the combination of the blank image (*B)*, the objects added (*O)* at position (x, y), and the noise following normal distribution (*N*).

The SNR of the stack is then estimated as,

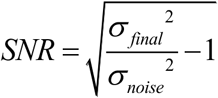

Where,*σ*_*final*_^2^ and *σ*_*noise*_^2^ are the variances of the final image stack and the noise, respectively.

While each frame in the single object datasets had one object, the multiple object datasets were constrained to include a total of 7 objects. Among these, 5 objects were consistently present from the first to the 148^th^ frame, while an additional object was introduced in the last two frames at different locations. This setup was designed to demonstrate the program’s capability to detect cells that appear later in the image stack.

### Experimental Methods

The imaging experiments shown in this manuscript are taken from previously published work where experimental details and procedures are described in detail [9, 22]. In brief, for the preparation of acute cortical or hippocampal tissue slices, BALB/c mice (both sexes) from postnatal day (P) 14-21 were anaesthetized, decapitated, and their brains removed. The brains were then placed in ice-cold preparation saline (in mM: 130 NaCl, 2.5 KCl, 0.5 CaCl_2_, 6 MgCl_2_, 1.25 NaH_2_PO_4_, 26 NaHCO_3_, and 10 glucose) saturated with 95% O_2_/5% CO_2_ to achieve a pH of 7.4. Subsequently, brains were cut into 250 μm thick slices using a vibratome (HM 650V, Thermo Fisher Scientific). For wide-field imaging, slices were kept for 20 min at 34°C in preparation saline containing the astrocyte-specific dye sulforhodamin 101 (SR101, 0.5 – 1 μM), followed by incubation for 10 min in artificial cerebrospinal fluid (ACSF, in mM: 130 NaCl, 2.5 KCl, 1.25 NaH_2_PO_4_, 26 NaHCO_3_, 2 CaCl_2_, 1 MgCl_2_, and 10 glucose, saturated with 95% O_2_/5% CO_2_, pH 7.4) at 34°C. Slices used for multiphoton imaging were kept at 34°C for 30 min in ACSF without SR101. Preparation saline and ACSF were both adjusted to an osmolarity of 310 mOsm l^-1^. Slices were kept dark in ACSF at room temperature (21°C) for up to 6 hours until use [22].

To measure changes in intracellular sodium ([Na^+^]_i_) or calcium ([Ca^2+^]_i_), slices were bolus loaded with the membrane-permeable form of the Na^+^-indicator ION-NaTRIUM-Green-2 (ING-2 AM; #2011F, Mobitec, Rheda-Wiedenbrück, Germany) or of the Ca^2+^-indicator Oregon Green 488 BAPTA-1 (OGB-1-AM; #O6807, Invitrogen, Waltham, Massachusetts, USA). Astrocytes were identified by selective labelling with SR101 [23]. High-resolution multiphoton imaging of ING-2 was performed using a custom-built laser scanning microscope based on a Fluoview300 system (EVIDENT Europe GmbH, Hamburg, Germany), equipped with a 60x water immersion objective (NIR Apo 60x, 1.0 W, Nikon Europe B.V., Amstelveen, The Netherlands). Excitation wavelength was 840 nm; fluorescence emission (filtered with 534/30, catalogue #F39-533, AHF Analysentechnik, Tübingen, Germany) was collected on a phosphomolybdic acid (PMA) hybrid photodetector (PicoQuant GmbH, Berlin) and registered by a MultiHarp 150 (PicoQuant). Pixel dwell time was 3.81 ms for frames of 512 x 512 pixels and at a typical frame rate of 1 Hz [22]. Wide-field imaging of OGB-1 was performed using an upright microscope (Eclipse FN1, Nikon) equipped with an ORCA FLASH 4.0LT camera (Hamamatsu Photonics Deutschland GmbH, Herrsching am Ammersee, Germany) and a 40x water immersion objective (Fluor 40×/0.8 W, Nikon). Probes were excited using a Polychrome V monochromator (ex: 488 nm, em: >505 nm; Thermo Fisher Scientific/FEI, Planegg, Germany) at a framerate of 1 Hz [9].

To induce intracellular ion signals, cellular metabolism was inhibited by perfusing slices with glucose-free ACSF containing 5 mM sodium azide (Riedel de Haen, Selze, Germany) and 2 mM 2-Deoxyglucose (Fluorochem, Hadfield, United Kingdom) for 2 minutes.

## Results

### Analysis of synthetically generated data

To assess whether the program can detect the correct number of fluorophore-loaded cells/ROIs, we utilized various sets of synthetically generated data. Figure 1A displays the generated data for multiple objects with their corresponding SNR values, accompanied by the true count of the cells. Each dataset was input into the program and subjected to pre-processing as needed to obtain segmented and labeled images. In addition to identifying cells that remained consistent throughout the stack, the program demonstrated its ability to detect cells appearing later in the image sequence. The cells identified by DL-SCAN are depicted in Figure 1B followed by the corresponding number of detected cells. While certain pre-processing steps aimed at reducing noise in images with low SNR values introduced a disparity between the actual and detected cell counts for some specific datasets (for example, SNR of 0.26), the program’s ability to identify cells from noisy background remained very good. In cases where a larger number of cells are detected, possibly influenced by the extent of preprocessing steps performed, users have the flexibility to selectively choose the cells of their interest and conduct analyses on those specific selections.

**Figure 1:**
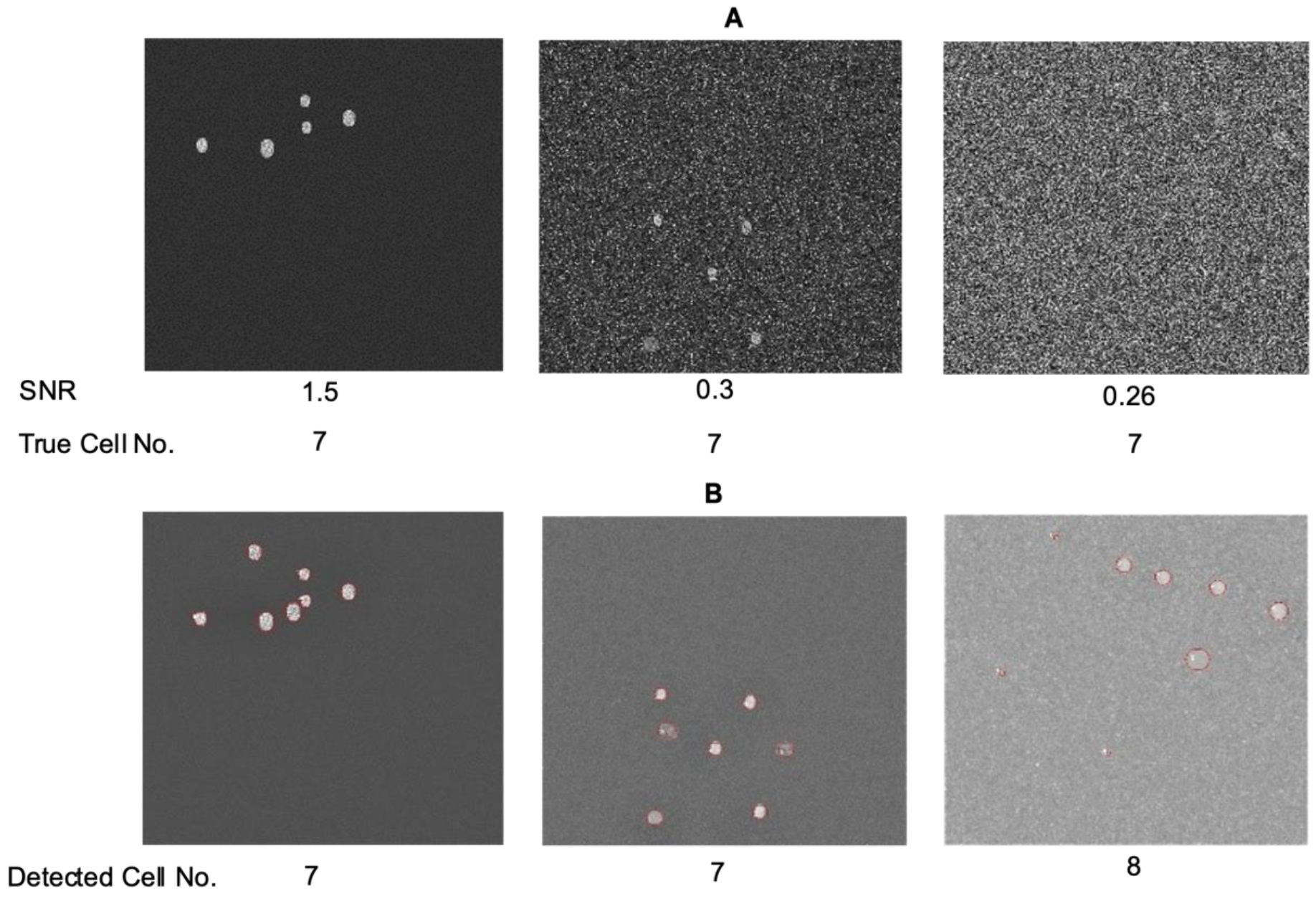
DL-SCAN can accurately detect cells in image stacks with different SNR values. First frames of synthetically generated images with a total of 7 cells (A) and DL-SCAN-identified cells on collapsed images (B) for varying SNR values.

To ensure that the program also detects the correct mean intensity as a function of time and associated parameters, we used distinct single-object datasets with a range of SNR values (Figure 2A). Correspondingly, Figure 2B displays the resultant segmented and labeled images. As observed previously, the program adeptly identifies the ROI in this scenario as well. The mean intensity traces generated by the program closely align with the true traces for all SNR values (Figure 2C).

**Figure 2:**
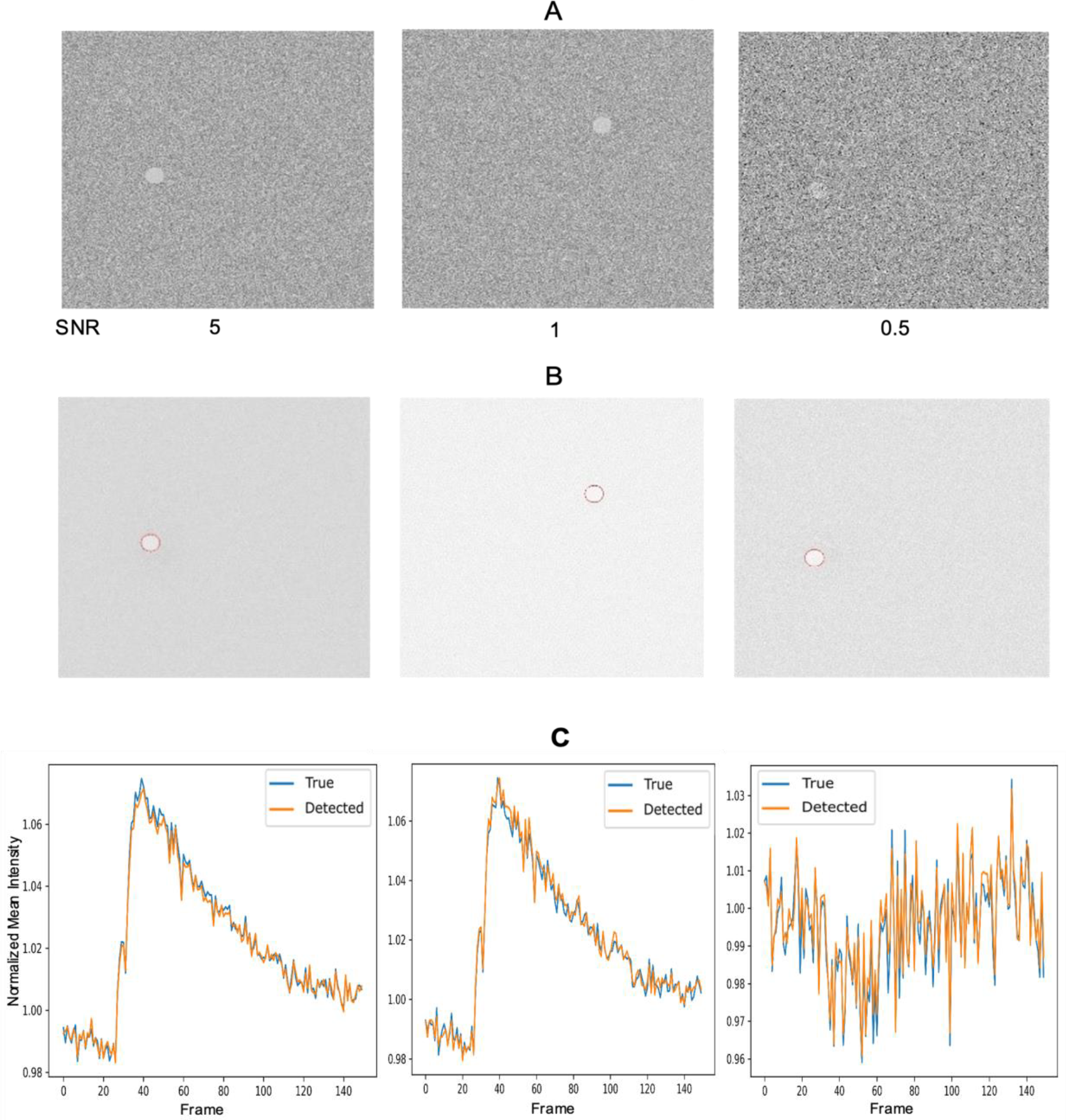
DL-SCAN can accurately extract time traces from images with different values of SNR. First frames of synthetically generated images with a single cell for varying SNR values (A) and DL-SCAN-identified cells on collapsed images (B). Comparison of the true normalized mean intensity with those extracted by DL-SCAN (C).

We also performed a comparison between the detected cell areas and their true areas with varying SNR, further reinforcing the effectiveness of the program, as shown in Figure 3. Although the program is also capable of computing the rise rate, rise time, decay time, and duration of the obtained trace for each selected cell, here we only illustrate the decay rate and amplitude for the given dataset for demonstration (Figure 3), which showcases the program’s remarkable efficacy. The experimentation conducted on the datasets using various additional options available in the program is outlined in the User Manual (Supplementary Information Text).

**Figure 3:**
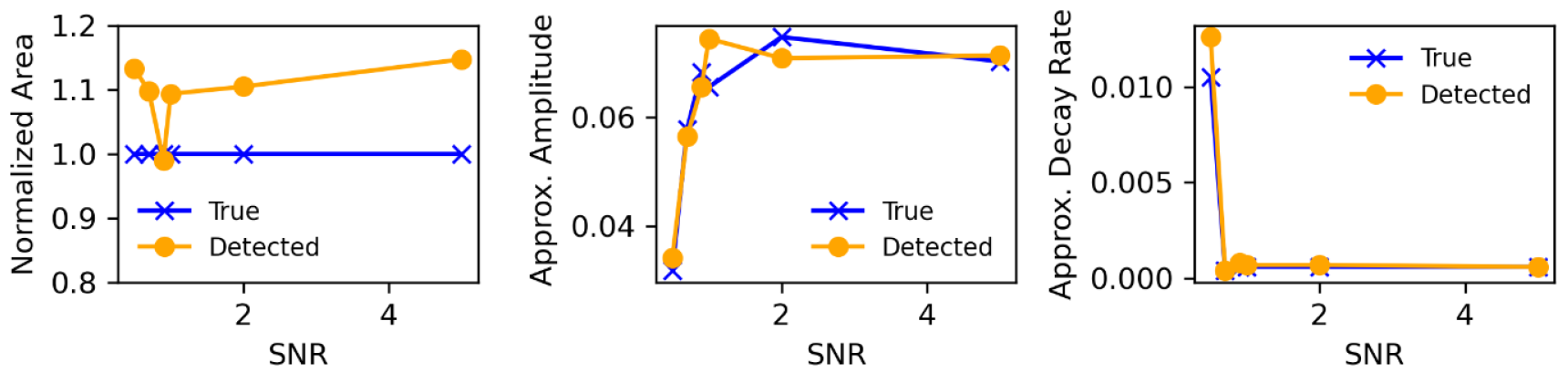
DL-SCAN can accurately extract related parameters from images with different values of SNR. Comparison of the true normalized area, amplitude, and decay rate (s^-1^) with those extracted by DL-SCAN.

### Application to experimental data

The next part of the validation process was implementing DL-SCAN on previously published experiments [9, 22]. In this work, acute tissue slices prepared from mouse brain were exposed to inhibitors of glycolysis and oxidative respiration for 2 minutes to induce chemical ischemia, resulting in transient increases in intracellular ion concentrations in neurons and astrocytes. Experimental data obtained from multiphoton imaging of ING-2 and wide-field imaging of OGB-1 for tracking intracellular Na^+^ and Ca^2+^ in hippocampal CA1 pyramidal neurons and layer II/III cortical astrocytes, respectively, were uploaded, segmented, and analyzed. The traces generated by using DL-SCAN were then compared to the manually analyzed results.

The traces generated by using DL-SCAN were then compared to the manually analyzed results. As shown in Figure 4, the changes in ING-2 fluorescence for the selected ROIs, representing neuronal cell bodies in the CA1 pyramidal cell layer, are almost identical. The minor discrepancies are most likely due to minimal differences in the ROI placement in the manual segmentation of the cells in the top panel of Figure 4A versus their automated identification as shown in Figure 4D. However, these variations are minor and not biologically relevant compared to the Signal-to-Noise Ratio in these experiments.

**Figure 4:**
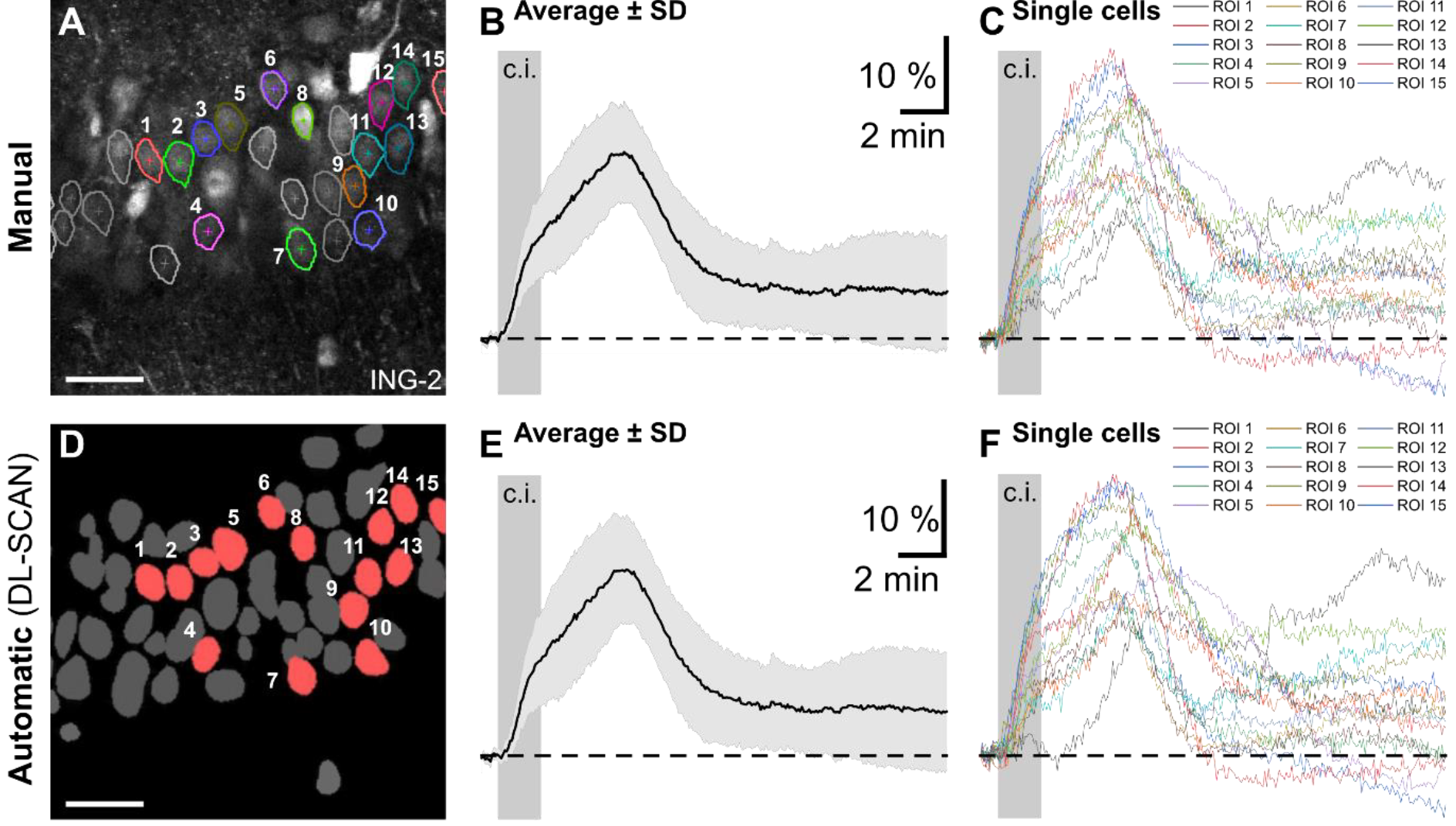
Automated segmentation and analysis by DL-SCAN of fluorescence microscopy imaging of changes in intracellular Na^+^ in hippocampal CA1 pyramidal neurons exposed to a 2-minutes period of chemical ischemia. Manual Segmentation using NIS Elements (A) and DL-SCAN-segmented neurons on collapsed image (D) with average (B and E) and individual (C and F) traces for the selected cells. The average intensity trace is represented by the solid line, with the standard deviation shown as error bars in light gray in both B and E. For accurate comparison, traces from the automated tool were obtained selecting “No Bleaching Correction” to be consistent with the published manual analysis. Scales 50 μm. Data taken from [22].

As a second proof-of-principle, we re-analyzed and compared changes in intracellular Ca^2+^ induced by chemical ischemia in astrocytes in cortical brain slices [9]. The changes in astrocytic OGB-1 fluorescence for manual and automatic segmentation are shown in Figure 5A and 5D. This part of the analysis was performed using the bleaching correction option. In line with the manual analysis published previously [9], the initial and final 10 frames were utilized for interpolating intermediate values. These values were subsequently subtracted from the original trace to correct for photobleaching. The normalized results obtained clearly demonstrate the close correspondence between the traces obtained from manual analysis and from DL-SCAN. The elimination of the negative values from the latter was done by subtracting the absolute minimum value from all intensity values. Although this explains the difference in the traces as the normalized intensity values drop below 0 in manual analysis, it has no significant effect on the subsequent analysis.

**Figure 5:**
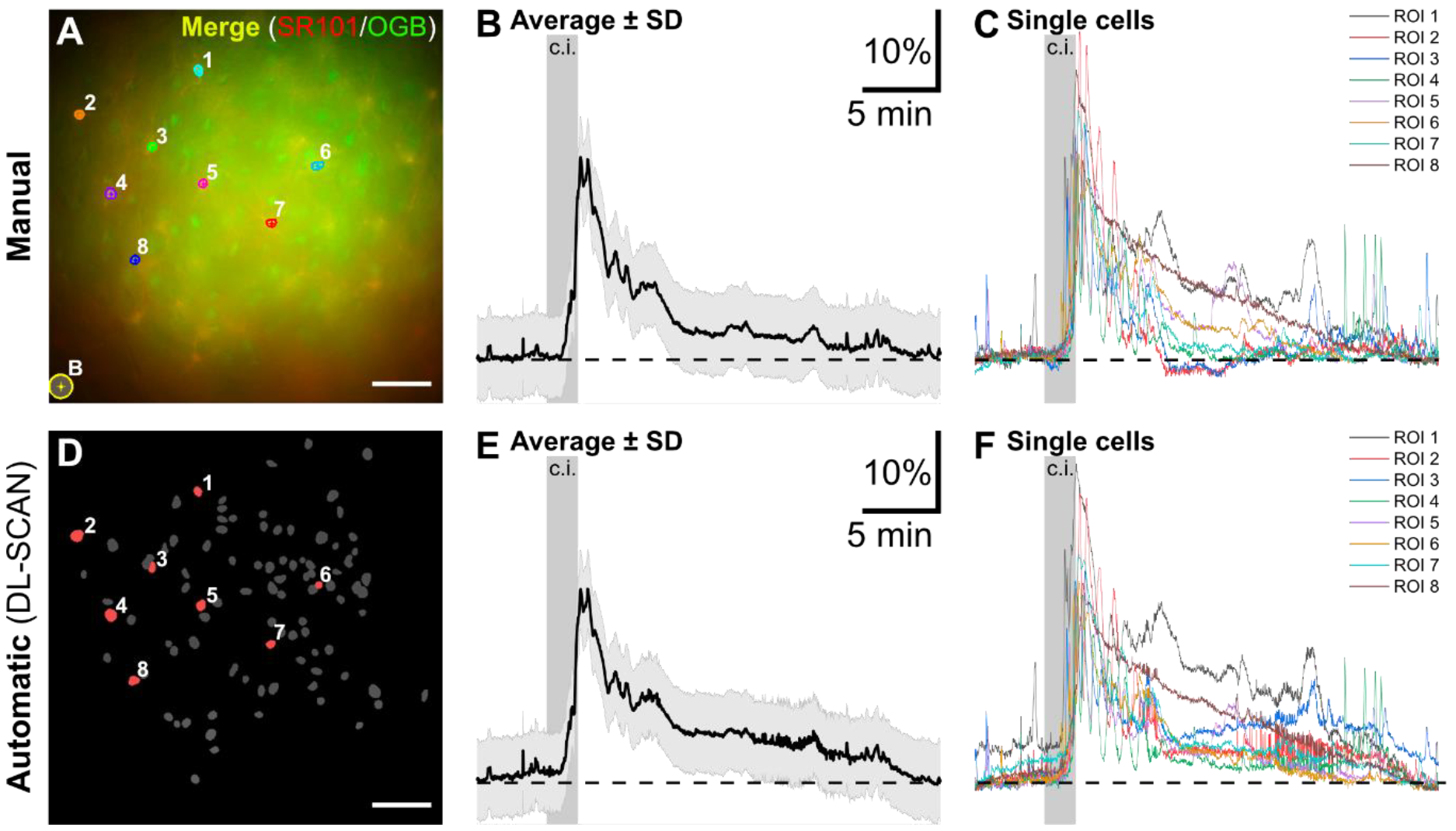
Automated segmentation and analysis by DL-SCAN of fluorescence microscopy imaging of changes in intracellular Ca^2+^ in layer II/IIII cortical astrocytes exposed to a 2-minutes period of chemical ischemia. Manual Segmentation using NIS Elements (A) and DL-SCAN-segmented astrocytes (D) with average (B and E) and individual traces (C and F) for the selected cells. The average intensity trace is represented by the solid line, with the standard deviation shown as error bars in light gray in both B and E.

The direct comparison between the manual and DL-SCAN analyses for each experiment (neuronal Na^+^ imaging and astrocytic Ca^2+^ imaging), shown in Figure 6A and B for two of the ROIs, also validates the efficacy of the program.

**Figure 6:**
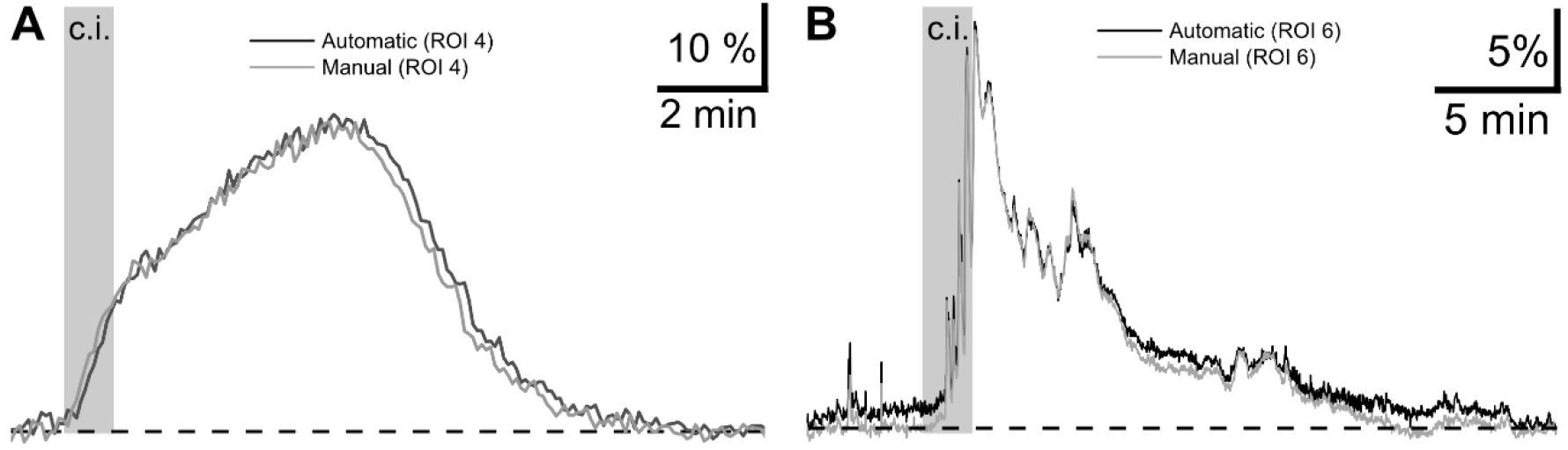
Time traces extracted by DL-SCAN show that the program accurately detects the temporal evolution of the signal. Direct comparison between manually and DL-SCAN-extracted traces (A) of changes in Na^+^ in two randomly selected hippocampal neurons and (B) in Ca^2+^ in two randomly selected cortical astrocytes exposed to brief chemical ischemia.

In Figure 7, we show DL-SCAN’s ability to generate various statistics for the selected ROIs from the segmented objects in another experiment in which changes in ING-2 fluorescence induced by chemical ischemia were measured in CA1 pyramidal neurons [22]. Contingent upon various post-processing options like baseline estimation, photobleaching correction, static or dynamic analysis etc., relevant information about rise time, rise rate, decay time, decay rate, duration, and amplitude of time traces can be generated for user-selected cells. These quantities provide instant, valuable data on the recorded signals in selected cells. For instance, the accumulation and clearance of ions in specific cellular compartments, as well as the comparison of the extent of ion fluxes across different experimental conditions or cell types, can be quantified using these parameters. DL-SCAN provides a very efficient and user-friendly way to perform such automated analysis. The statistics for the selected neurons that are shown in Figure 7 were generated by selecting “Dynamic” analysis option, with the baseline estimated as the average of the first 10 frames. The program provides bar plots of these parameters for each cell (Figure 7). The relevant data can also be exported in ascii format for further analysis and visualization.

**Figure 7:**
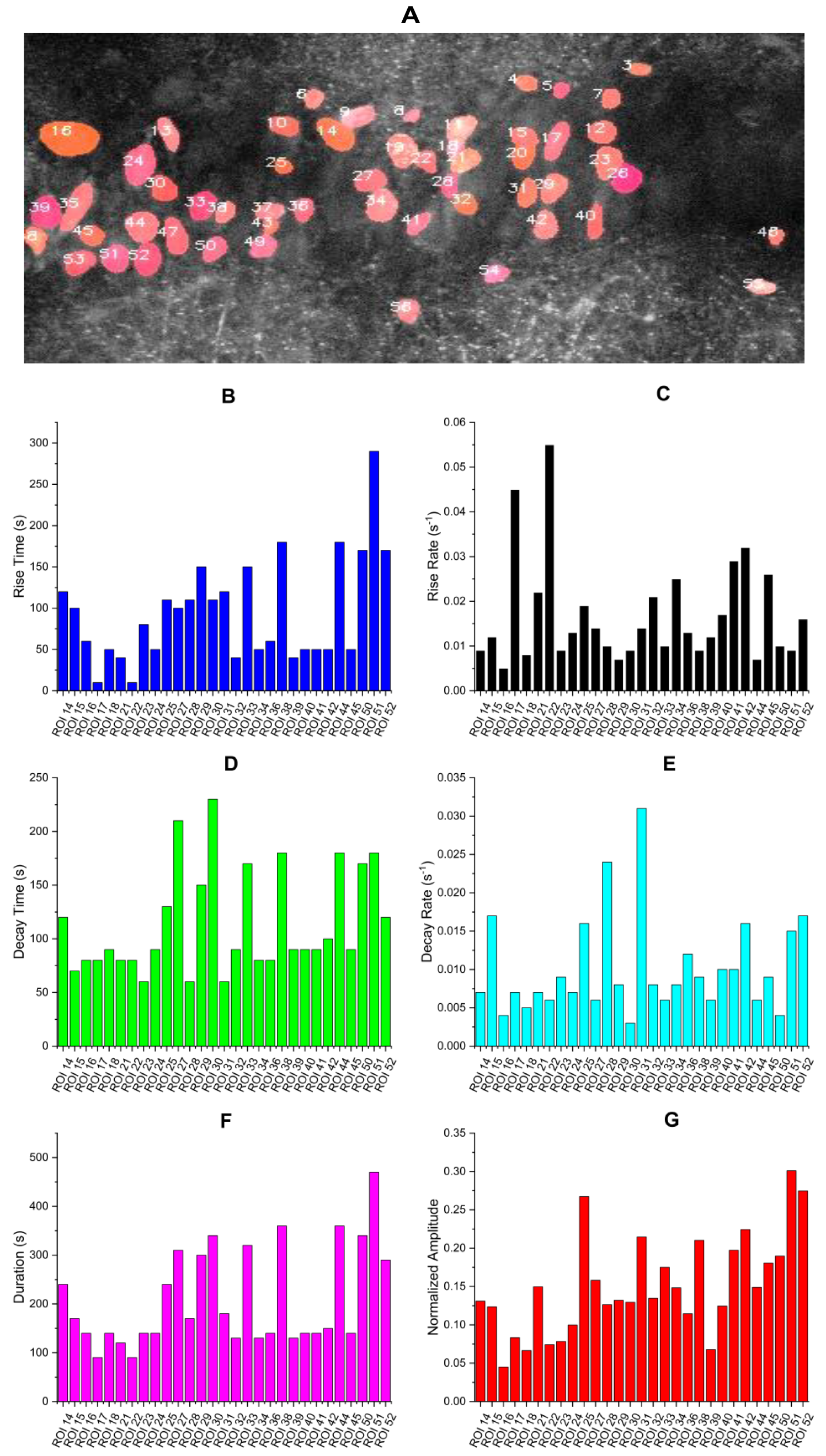
DL-SCAN provides analysis of relevant parameters on a cell-by-cell basis. (A) Automatically segmented and labeled neurons in an ING-1 loaded hippocampal tissue slice. Bar plots show properties of the transient changes in intracellular fluorescence upon brief chemical ischemia: rise time (B, Mean ± SD, 95 ± 63 s), rise rate (C, 0.017 ± 0.012 s^-1^), decay time (D, 114 ± 48 s), decay rate (E, 0.010 ± 0.006 s^-1^), duration (F, 209 ± 101 s), and normalized amplitude (G, 0.15 ± 0.06) for the selected cells using DL-SCAN. Each bar in panels B-F corresponds to a single cell.

As mentioned above, the program also generates histograms of decay rates, rise rates, decay times, rise times, durations, and normalized amplitudes of traces from all cells detected in the image stack. In Figure 8, we show these distributions for cells detected in Figure 7A.

**Figure 8:**
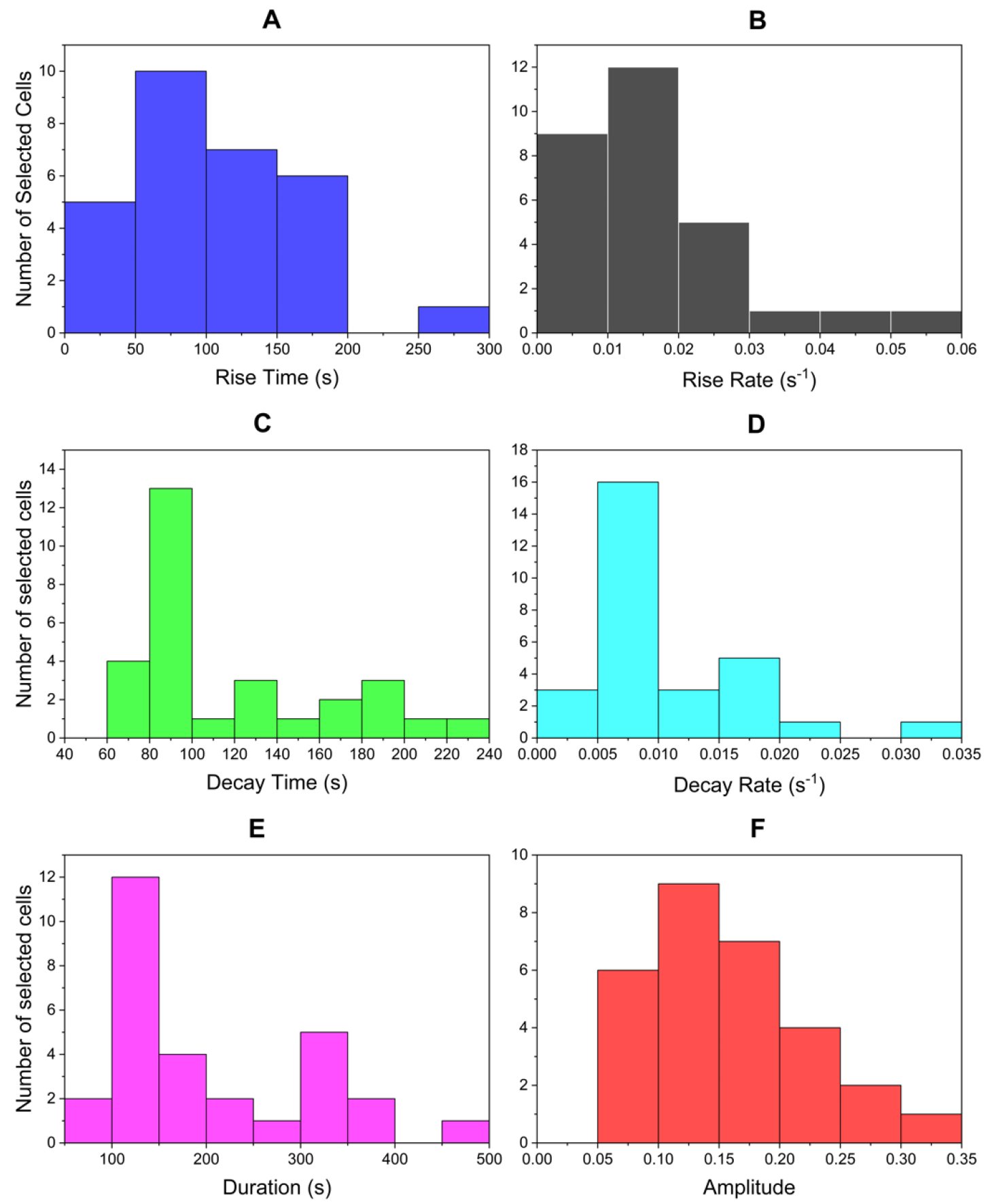
DL-SCAN can also generate histograms of various parameters for all cells detected. Distributions of DL-SCAN-generated rise time (A), rise rate (B), decay time (C), decay rate (D), duration (E), and normalized amplitude (F) of traces from the selected cells.

### Quantification of morphological changes in cells

Next, we demonstrate the tool’s ability to account for the morphological changes in cells, with the help of synthetic data. Preferably, quantifying cell swelling or shrinking requires a specific cell-permeant dye like Calcein-AM that diffuses across cytosol and remains inside the cell for longer period, emitting green light under blue light excitation [24, 25]. The longer retention time then becomes useful in monitoring the changes in cell’s area over time. To demonstrate this ability, we employ synthetic data generated to represent swelling in a cell stained with a Calcein-like dye by gradually increasing the size of the object and decreasing it again to normal over time. The corresponding results are shown in Figure 9. The counts of bright pixels (pixels exceeding the threshold multiplied by the maximum pixel value) are obtained using the default area percentage threshold of 0.3. This essentially refers to the count of pixels within each ROI that exceed 30% of the maximum pixel value. Figure 9 effectively estimates the change in the morphology of cells over time when segmented on the collapsed image, demonstrating the ability of DL-SCAN to track changes in cell’s area.

**Figure 9:**
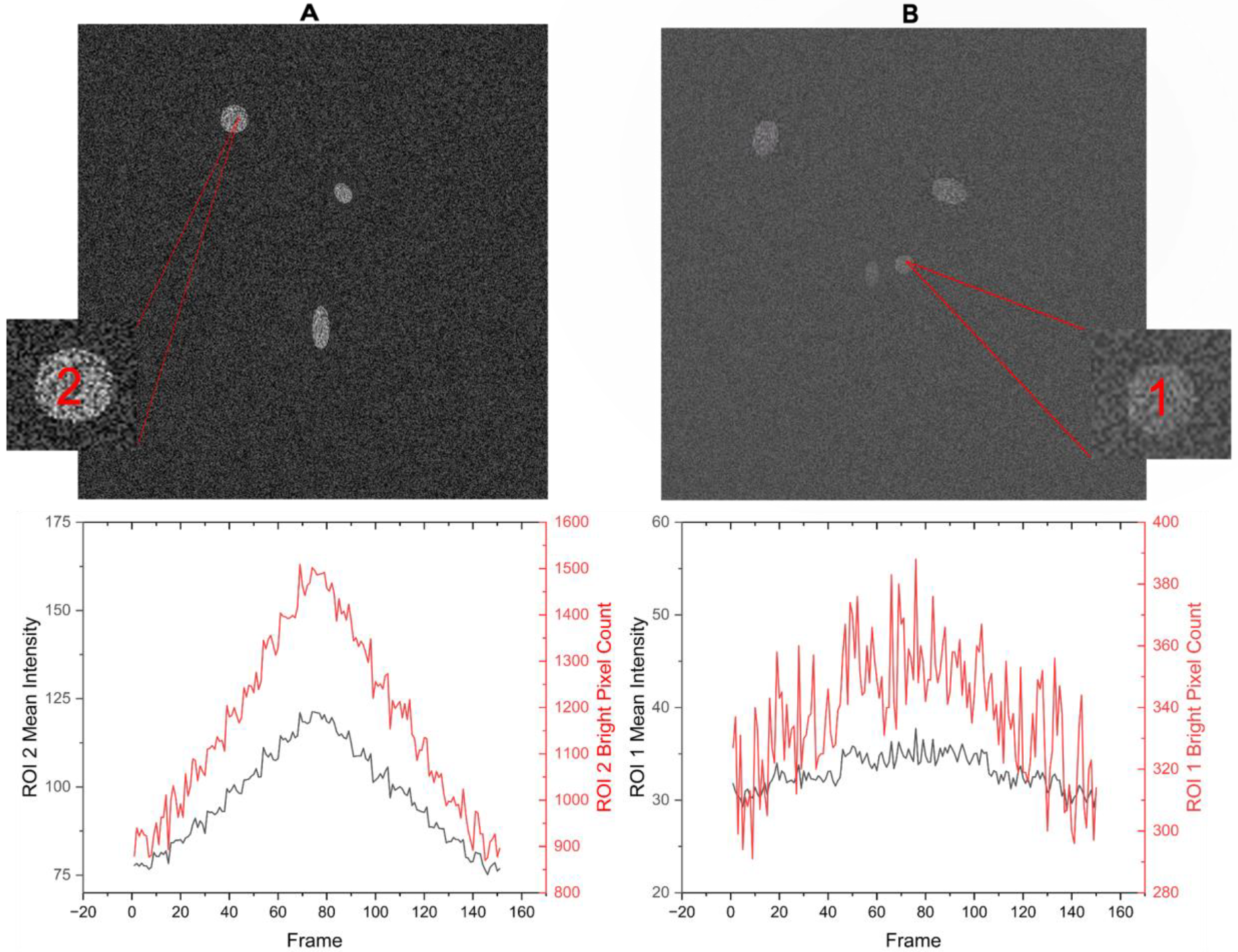
DL-SCAN can track the temporal changes in the area of cells in the image stack. DL-SCAN segmented synthetically generated cells (A and B) (upper panels). Examples of the change in the cell’s area (bright pixel count) and mean intensity for the generated synthetic data (A and B) (lower panels).

## Discussion

The existing microscope manufacturer’s tools like NIS Elements by Nikon Instruments or CellSense FluoView by Evident Europe primarily require manual selection of ROIs for analysis. The reliance on this manual labeling and analysis is time-consuming and subjective, particularly when working with large datasets or complex images. Automated tools like AQuA (Astrocyte Quantification and Analysis), CaSCaDe (Ca^2+^ Signal Classification and Decoding), and MSparkles identify ROIs based on changes in intensity of neighboring pixels [14-16]. Instead of picking specific regions to study beforehand, AQuA uses the whole image to find patterns in how fluorescence changes over time, while MSparkles uses a region-growing approach to identify ROIs [14, 15]. While tools like these offer flexibilities in terms of algorithm customization and parameter tuning, which can be beneficial for researchers with specific needs or for analyzing diverse datasets, Convolutional Neural Network (CNN)-based approaches prove to be very effective in terms of accuracy even for generalizing variable image patterns and tasks like ROI identification. The various components of a CNN architecture, such as convolutional layers, pooling layers, skip connections, regularization layers, loss functions and activation functions, work together to extract and preserve complex features, spatial information, and details from the input image in an unprecedented manner.

To overcome the shortcomings in the existing approaches mentioned above, we developed DL-SCAN, which is user-friendly, yet equally effective for cell segmentation and analysis. The higher Average Precision (AP) cell segmentation accuracy of StarDist as compared to other popular object segmentation models for various intersection over union threshold values proves to be a significant advancement in the field of automated cell segmentation [17]. Its performance showcases its robustness and accuracy in delineating cell boundaries, which is crucial for biological applications. Accurate cell identification immensely helps in correct quantification of cellular features, cellular functions, cellular localization, and cellular interactions. It is also essential for visualizing and analyzing cells in tissues, which is critical for identifying and monitoring the diseased state. Therefore, we decided to incorporate StarDist into our tool, resulting in high-quality cell segmentation outcomes. This is followed by various user-adjustable options to facilitate and streamline efficient analysis as needed by generating relevant statistics. Although DL-SCAN generates publication-quality figures incorporating a comprehensive analysis of the data, options are provided to download the generated files in CSV format to enable users to upload the data to external software for further processing, analysis, and visualization. We believe that this tool will benefit researchers and experimentalists in the imaging field to study cells and/or cellular compartments by significantly reducing time consumption and bias, while increasing accuracy.

Synthetic data with varying SNR, which were representative of cellular somata (neurons and astrocytes), were generated to test the accuracy of the cell segmentation and their statistics. Secondly, the tool was tested on Na^+^ imaging data from hippocampal neurons and Ca^2+^ imaging data from cortical astrocytes under chemical ischemia published previously [9, 22]. The outcomes were then compared to the previously published analyses performed using NIS Elements software from Nikon instruments [9, 22]. As opposed to the time-consuming manual analysis performed before, DL-SCAN easily and rapidly managed to automatically segment somatic ROIs. Furthermore, DL-SCAN enabled immediate extraction of their fluorescence signals over time as well as of other parameters, also offering numerous user-adjustable options for pre-processing, and options related to the specific experimental setup. These features help to delineate and objectify the entire image analysis process.

Along with intensity changes over time, this tool also records bright pixel counts above a certain user-specified threshold for each detected cell. Since cells can be segmented based on the first or the collapsed image, the “Area Threshold Percentage” option is introduced to account for cell swelling when segmentation is performed on the latter. The pixel count of a region greater than the threshold percentage of the maximum pixel value for that region is generated and plotted. This helps to track the changes in the area of the identified cell over time, as illustrated earlier, but surely has some limitations. This feature of the tool can, for example, be used with a fluorescent indicator that is inert to changes in ion concentrations and does not respond to other cellular signals,. For dyes that get brighter when they are binding to ions as the signal spreads spatially inside the cell (e.g., the propagation of a Ca^2+^ wave in the cytoplasm), recording the pixel count surpassing a certain brightness threshold may not work. Nevertheless, the algorithm will accurately track changes in cell area in images where the ROI is changing due to cell swelling.

To summarize, the major benefit of DL-SCAN is its ease of use with a wide variety of datasets, its accuracy, avoiding user-bias as the Deep Learning algorithm automatically segments and analyzes the cells, and its access to additional features and statistics that are not incorporated in other tools. Although tailored for a certain use case (acute brain slices), the nature of the detection algorithm and the subsequent analysis options, together, makes it easily adaptable for a broad range of applications. Furthermore, our open-source algorithm provides an opportunity to update and customize the application as needed.

## Supporting information

Supplemental Text (User Manual)

## Acknowledgements

This work was supported by National Institutes of Health through grant number R01NS130916 (to G.U.), by the German Research Foundation (DFG), Research Unit 2795 “Synapses under stress” (Ro2327/13-2 to C.R.R.) and a Mercator fellowship (to G.U.) and by the Federal Ministry of Education and Research (BMBF), Germany (Project SynGluCross to C.R.R.).

