## Supplemental Text (User Manual) for "A Deep Learning-Based Segmentation of Cells and Analysis (DL-SCAN)"

DL-SCAN, which effectively segments and analyzes cells in images from fluorescent microscopy (in TIFF format), is developed using Streamlit library (version 1.21.0) in Python 3.8.8. The user-friendly graphical user interface (GUI) displayed immediately after the tool is launched is as shown (Figure S1).

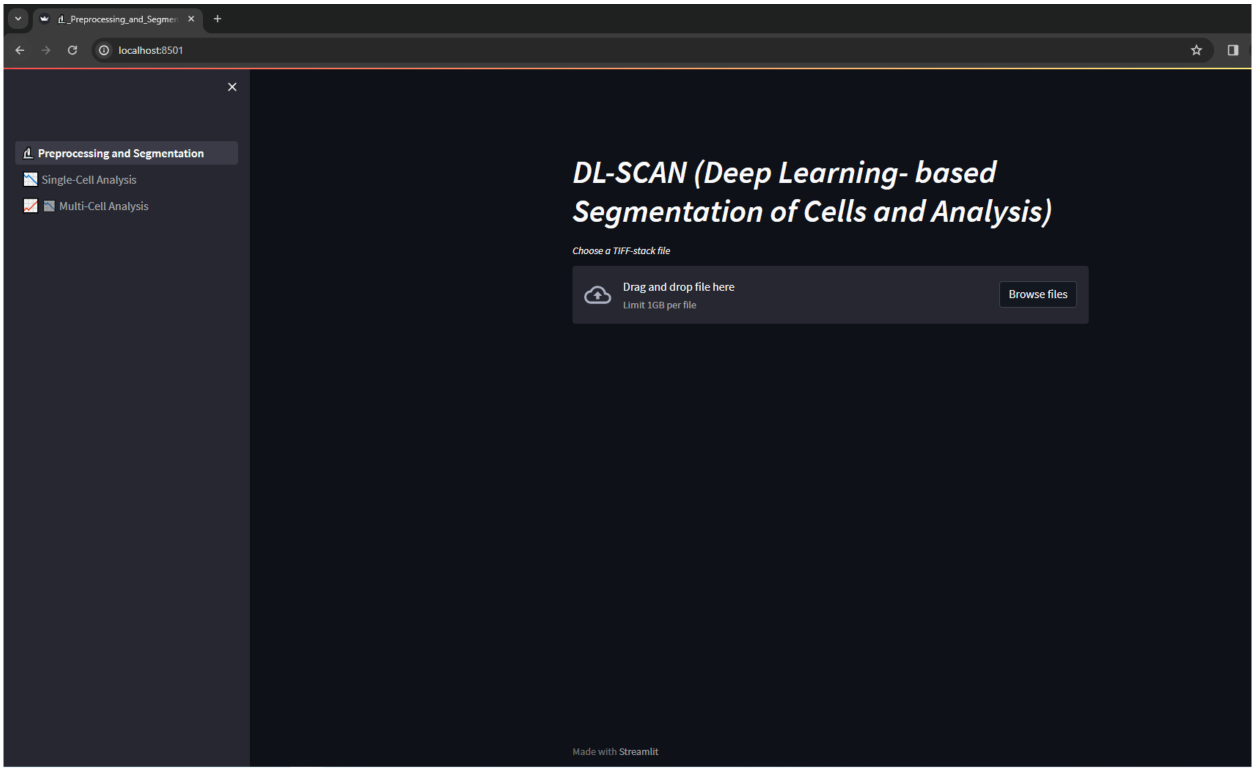

**Figure S1.** Graphical User Interface of DL-SCAN in the browser. It consists of three main sections in the sidebar for preprocessing and segmenting images, single-cell analysis, and multi-cell analysis. The homepage (Preprocessing and Segmentation) asks the user to upload a TIFF image stack for preprocessing and segmentation.

### A. Launching the Application

**Note:** DL-SCAN is also available online on Hugging Face Spaces. This web application can be accessed from: https://huggingface.co/spaces/dlscan/dlscan.

For larger files, faster performance, and customization, running DL-SCAN locally is recommended. Using the Anaconda distribution is recommended for setting up and running the program. Link: **https://www.anaconda.com/download**

Since Streamlit, by default, provides only 200 MB free space for deploying and using Applications, a configuration file is written to increase the limit up to 1 GB, which can again be expanded further as shown later in the following steps.

1. Clone the provided GitHub repository to the local machine.
   https://github.com/banalok/dlscan

2. Open the Anaconda Command Prompt and navigate to the cloned repository destination. Create a new Python environment called *dlscan*, and install all the dependencies specified in the "requirements.txt" file using the following commands.

   *conda create -n dlscan python==3.8.8*

   *conda activate dlscan*

   *pip install -r requirements.txt*

3. Once all the dependencies are installed, the users can always execute the program by navigating to the directory where the Application is located (if not already in the directory), and activating the environment (if not already activated), and launch DL-SCAN by entering the following commands.

   **Option 1 (if the dataset is less than 1 GB)**

   *streamlit run DL_SCAN.py*

   **Option 2 (if the dataset is larger than 1 GB)**

   By default, the maximum upload size is 1 GB, but can be adjusted by adding " --server.maxUploadSize 2000" at the end of the command line, where the number 2000 represents the size in MB.

   *streamlit run DL_SCAN.py –server.maxUploadSize 2000*

   This size can also be adjusted in "config.toml" file inside ".streamlit" folder.

### B. Uploading Files

A microscopy image stack in 8-bit TIFF format can be uploaded by clicking "Browse" on the homepage of the program. The original frame of the uploaded file is displayed. This is followed by the option to incorporate background correction (Figure S2).

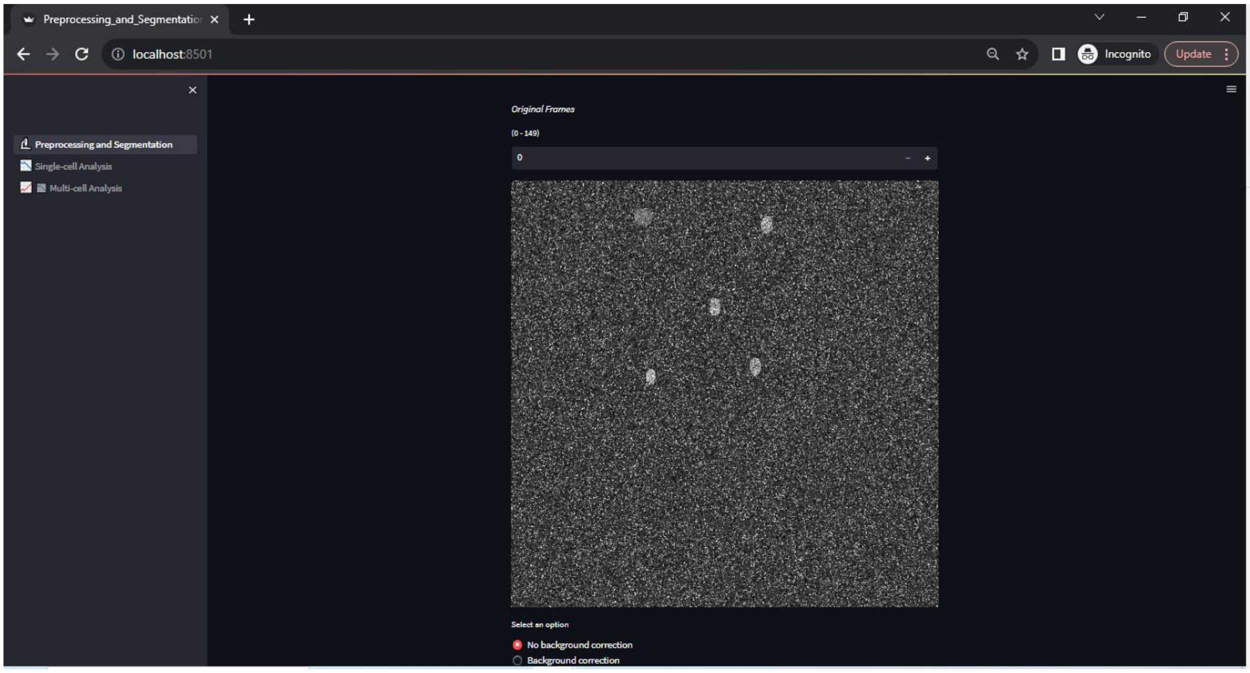

**Figure S2.** The first image frame out of 150 frames from the uploaded TIFF stack, followed by the option to incorporate background correction. Here, "No background Correction" is selected.

In case the user wishes to discard the current file and upload a new one, a simple page refresh will let them do that.

### C. Preprocessing options

#### Background Correction

When background correction is required, users can select "Background Correction" option. This action will display the first frame, allowing them to draw a rectangle on it (Figure S3-A). The average intensity of the selected pixels is then calculated and subtracted from the original image in each frame. The background corrected frames are then displayed (Figure S3-B). Users can input the frame number or click "+/-" to view the resulting background-corrected images.

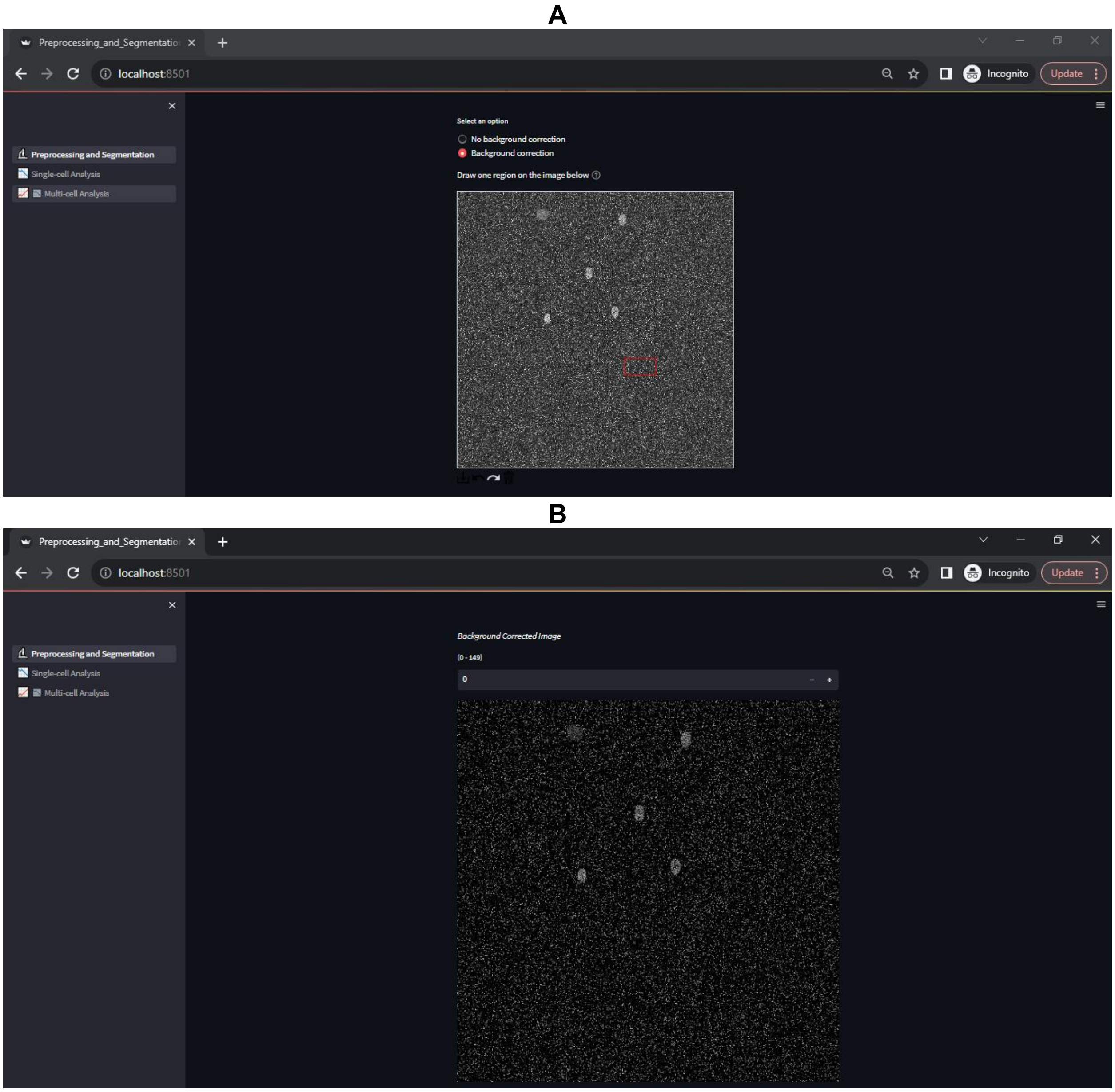

**Figure S3.** Users are presented with an option to draw a rectangle on top of the first image (A). The background corrected frame as a result of the drawing is displayed in (B).

### Gaussian Blurring

The first preprocessing option provided in the tool is Gaussian Blurring. The Gaussian square matrix generated as a result of the selected kernel size convolves the original image, calculating the weighted average to replace its center pixel value. This weighted averaging smooths the image (Figure S4), as the pixels closer to the center of the kernel have higher weights as compared to those that are far away.

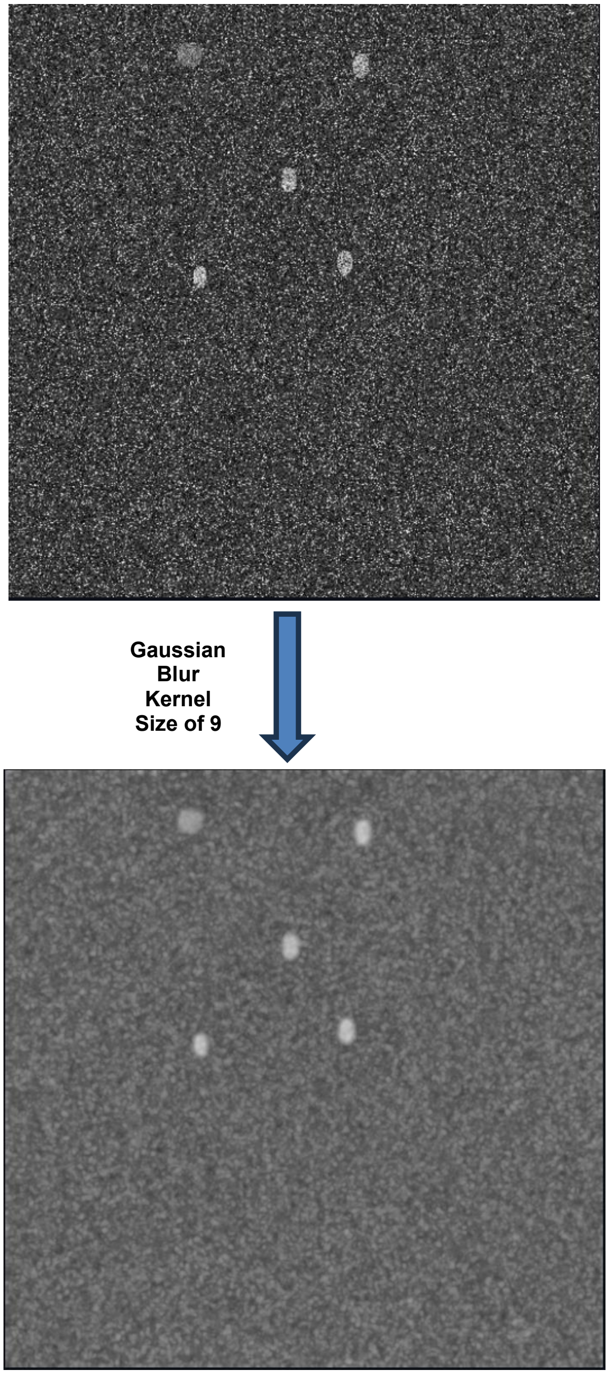

**Figure S4.** Gaussian Blur of kernel size 9 applied to the original image, improving the image quality by reducing noise, while preserving edges.

### Median Blurring

Users have the option to use median filtering by providing the Median Blurring Kernel Size. This process generates a square matrix of the selected size that slides on top of the image, computing median of the window and replacing the center pixel value. This works for reducing noise in an image, preserving edges and sharp features of the objects (Figure S5). More specifically, salt-and-pepper noise can be well-reduced using this technique.

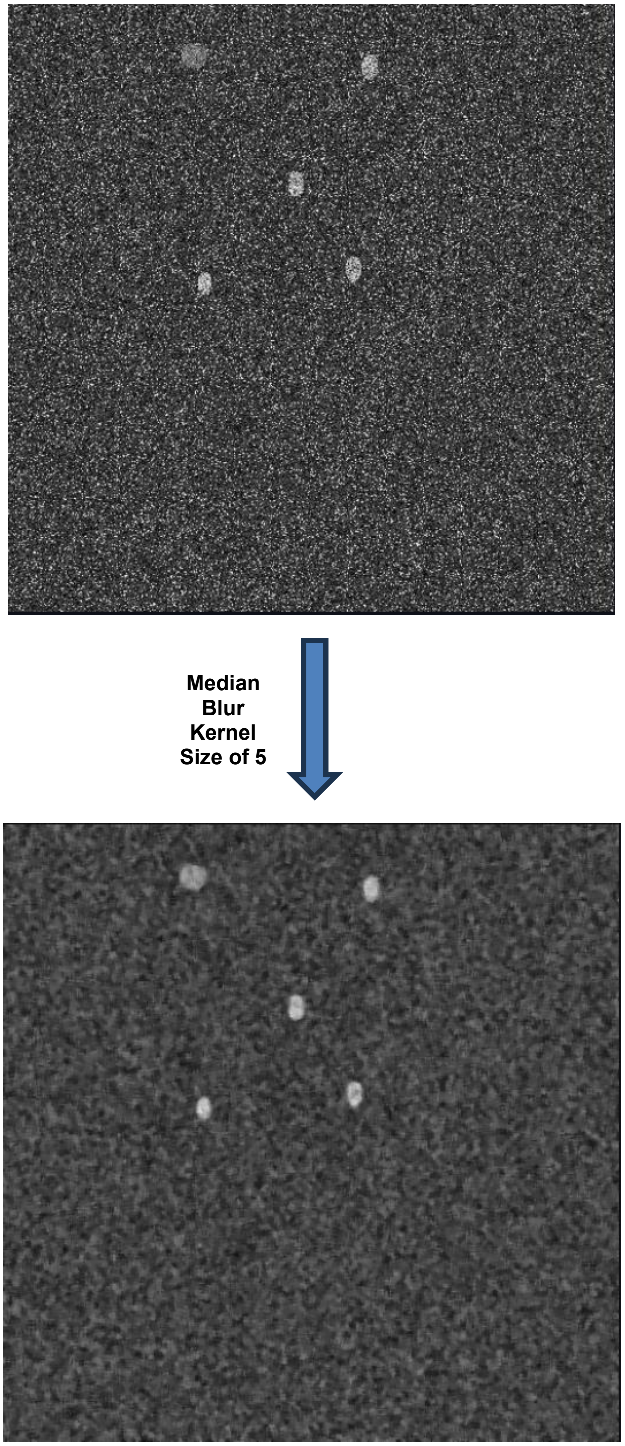

**Figure S5.** Median Blur of kernel size 5 applied to the original image, improving the image quality by reducing noise, while preserving edges.

### Brightness and Contrast

Image brightness and contrast can also be adjusted in the program as one of the preprocessing options.

### Contrast Limited Adaptive Histogram Equalization (CLAHE)

To prevent over-amplification of noise, Contrast Limited Histogram Equalization (CLAHE) is provided as one of the options in the program. Unlike traditional histogram equalization, CLAHE adapts the contrast enhancement locally within smaller regions of the image ($8 \times 8$ grid). Adjusting the provided clip limit factor value changes the amount of contrast enhancement (Figure S6). Although the higher value allows for more enhancement, one must be cautious using this option, as it can also lead to a noisier result.

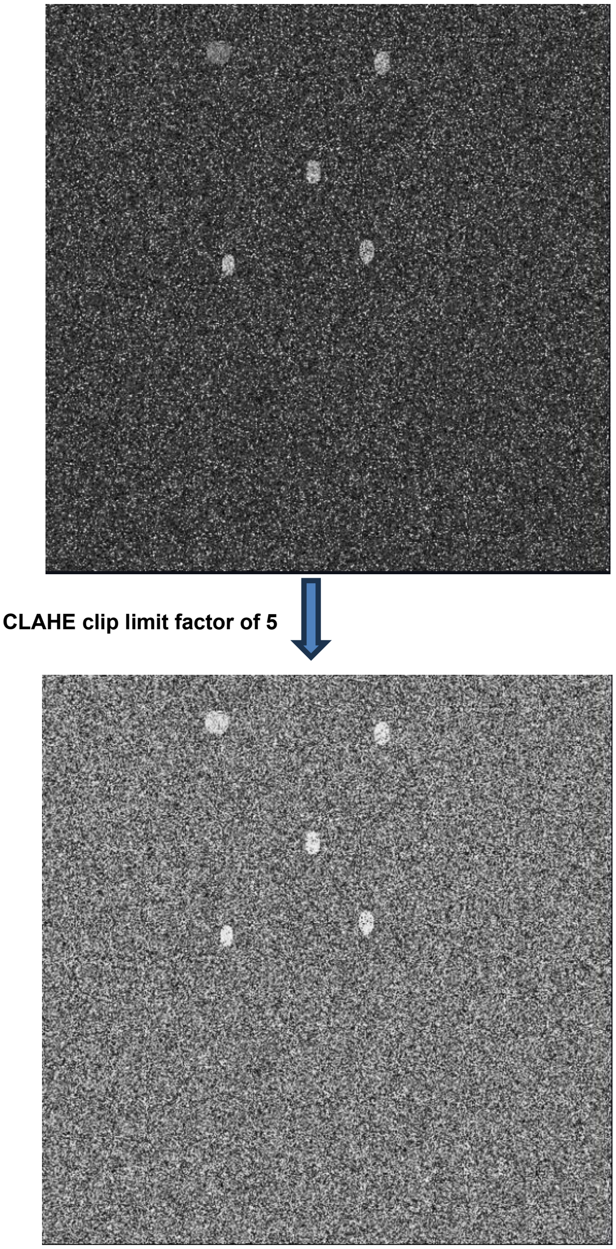

**Figure S6.** CLAHE clip limit factor of 5 applied to the original image, improving the image quality by bringing out details and features in both dark and bright regions of an image.

Users are encouraged to experiment with individual options, examining the outcome before applying all options simultaneously. This approach ensures effective image preprocessing.

### Processed Frames

The preprocessed frames, as a result of any of the applied preprocessing options or the combination of them is displayed as “Processed Frames”. Users can input the frame number or click “+/-” to view the resulting processed images.

### Collapsed Image

To account for cells appearing in subsequent frames, the entire stack of images is collapsed into a single image. In this process, pixels with higher values are retained during pixel-by-pixel comparison between frames (Figure S7). Therefore, it is crucial to upload a TIFF stack where cells are represented by higher pixel values and the background by lower pixel values.

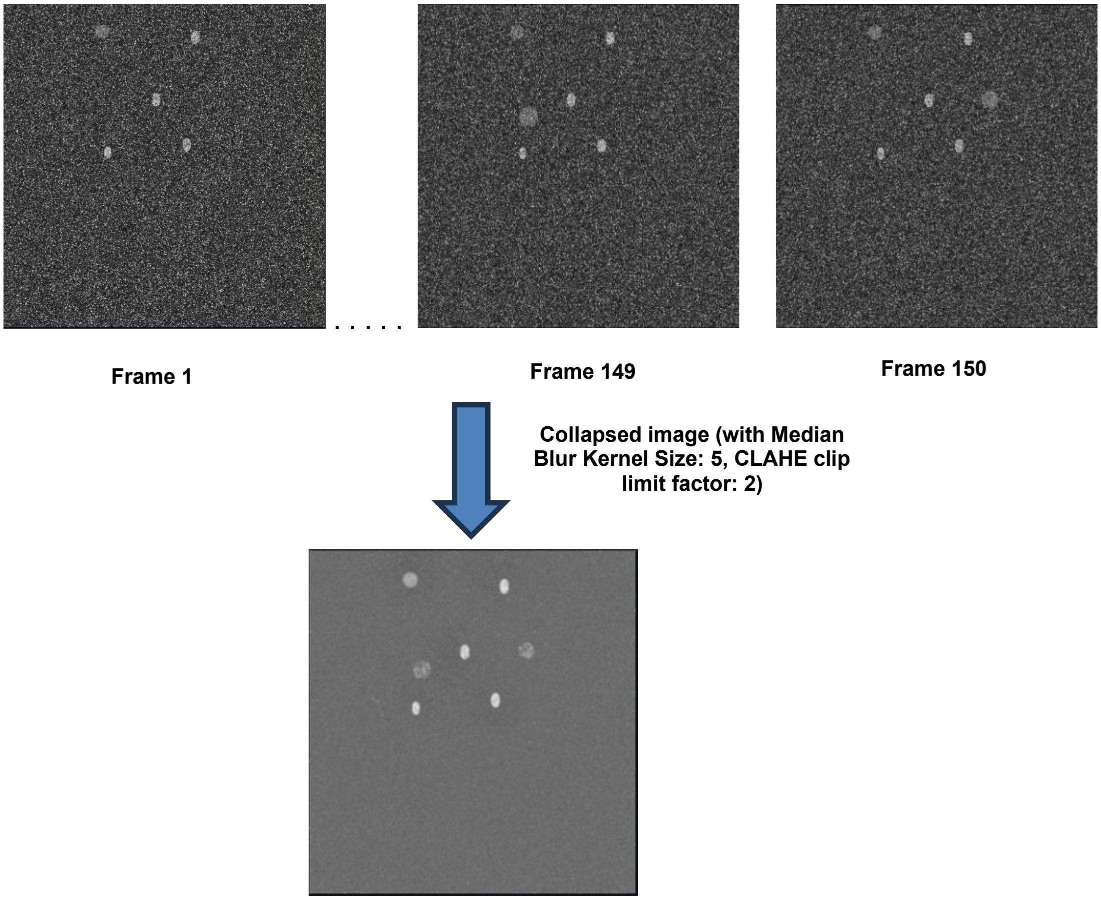

**Figure S7.** The original TIFF-stack with 150 frames, where the last two frames have an additional cell. The collapsed image accounts for all the cells throughout the stack and is ready for segmentation.

### D. Segmentation

The collapsed image is now ready for segmentation. Clicking “Segment and generate labels” outputs a segmented image and labeled image (Figure S8) and is ready for analysis.

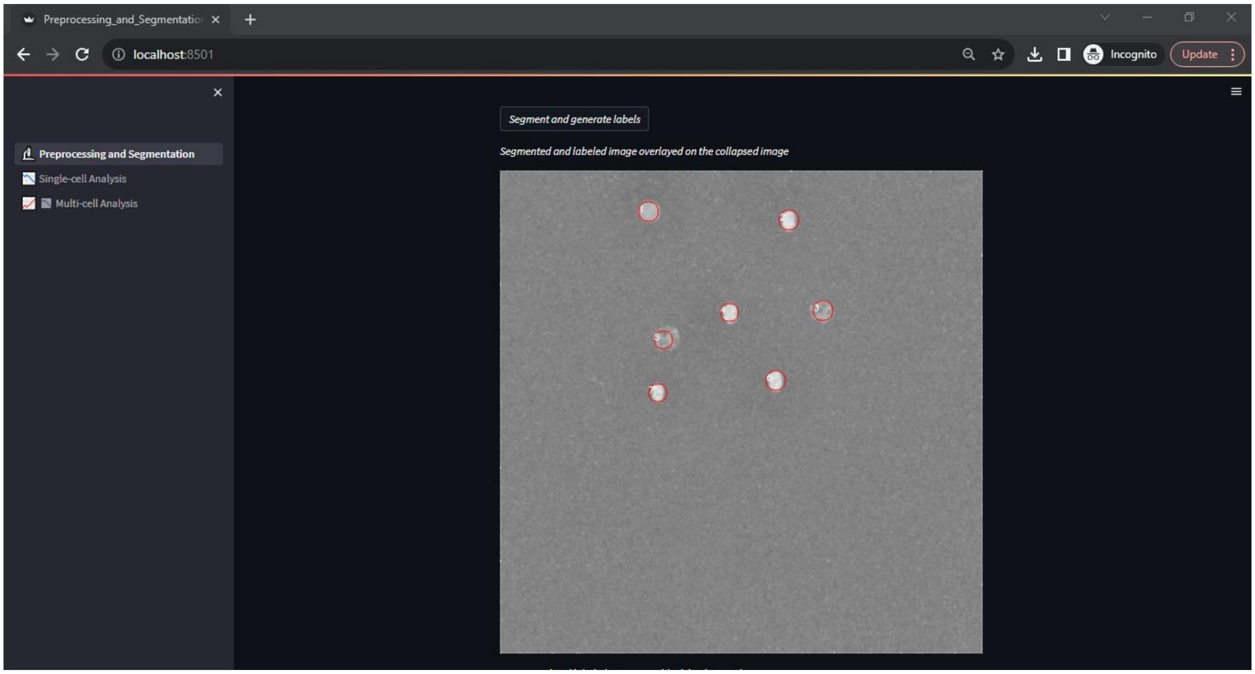

**Figure S8.** The segmented and labeled regions on top of the collapsed image after the segmentation algorithm is applied. The red circles do not represent the area of the segmented cells; they are solely used for numbering purposes.

At times, the pixel distribution in uploaded images can be uneven due to differences in the experimental setup, resulting in overexposure in some areas and underexposure in others. To address this, we offer a **Rolling Ball Background Correction (RBBC)** option, which helps mitigate this problem. A ball of a user-selected radius rolls across the image, fitting into its valleys and crevices. The minimum pixel value within the region covered by the ball is subtracted from the pixel value at the center of the ball, effectively smoothing out the background when unevenly illuminated.

Additionally, users also have the choice to segment based on the first image rather than the collapsed image.

In case needed, users have an option to manually draw regions of interest (ROI) that get added to the list of cells to be analyzed (Figure S9). Once segmented, a dye-positive image can be uploaded and overlaid on top of the segmented image for efficient cell selection as needed.

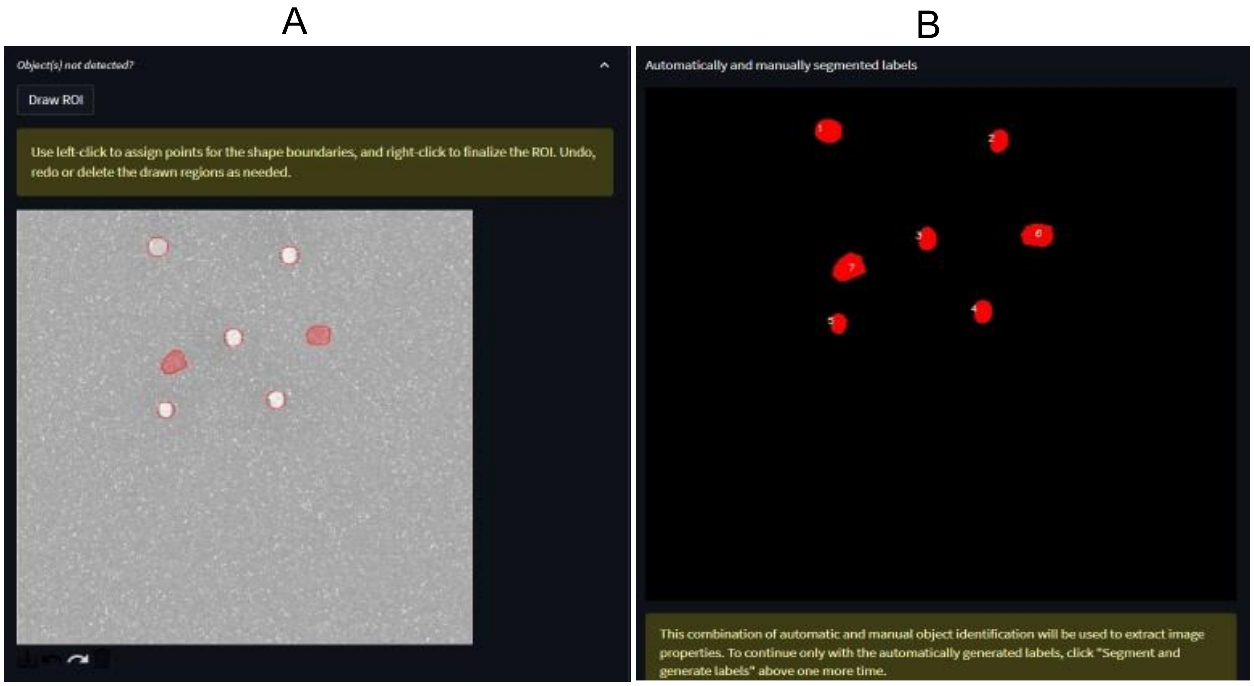

**Figure S9**. Manual ROI selection. (A) Two regions are drawn manually as required. (B) The drawn regions are displayed as additional labels on the automatically labeled image.

### E. Single-cell Analysis

This page can be accessed by clicking "Single-cell Analysis" in the sidebar. In this section, users can perform analysis of individual cells. To access the content of the page, the program assumes that the Preprocessing and Segmentation steps have been already completed. After displaying the collapsed and labeled images for reference at the top, a table is presented, followed by the same table, but now interactive. The first table can be downloaded as a CSV file and has the following format.

| label | intensity_mean_0 | intensity_mean_1 | ...... | intensity_mean_n | Bright_pixel_area_0 | Bright_pixel_area_1 | ...... | Bright_pixel_area_n |
|---|---|---|---|---|---|---|---|---|
| 1 | mean_1_0 | mean_1_1 | ...... | mean_1_n | area_1_0 | area_1_1 | ...... | area_1_n |
| 2 | mean_2_0 | mean_2_1 | ...... | mean_2_n | area_2_0 | area_2_1 | ...... | area_2_n |
| ...... | ...... | ...... | ...... | ...... | ...... | ...... | ...... | ...... |
| m | mean_m_0 | mean_m_1 | ...... | mean_m_n | area_m_0 | area_m_1 | ...... | area_m_n |

Here, the number of frames ranges from 0 to n and the number of cells (labels) from 1 to m. The "Bright_pixel_area" column corresponds to the number of pixels that are higher than a user-selected threshold (default is 0.3) provided by "Choose the area threshold percentage" widget. The interactive table allows users to select a single cell and perform analysis on it. As soon as the cell is selected, it is isolated and shown in the image highlighted by red color. This is followed by the option to input frame rate and the bleaching correction option with "No bleaching correction" option as the default.

### No bleaching correction

When this option is selected, users will get further options to

1. Adjust the moving average window for trace smoothing, that ranges from 1 to 5 (where 1 would mean the original trace) (Figure S10-B).
2. Select “Static” or “Dynamic” analysis to be performed (Figure S10-B).

**Static Analysis** lets users select a single baseline intensity frame number (“Single Frame Value”) or number of consecutive frames to average their intensity values for baseline intensity calculation (“Average Frame Value”), peak intensity frame number and recovery intensity frame number.

**Dynamic Analysis** asks users to select number of consecutive frames to average their intensity values for baseline intensity calculation, while the peak and recovery are automatically computed.

The normalized intensity table is then displayed, followed by the original, smoothed, and bright pixel area traces (Figure S10-C).

A

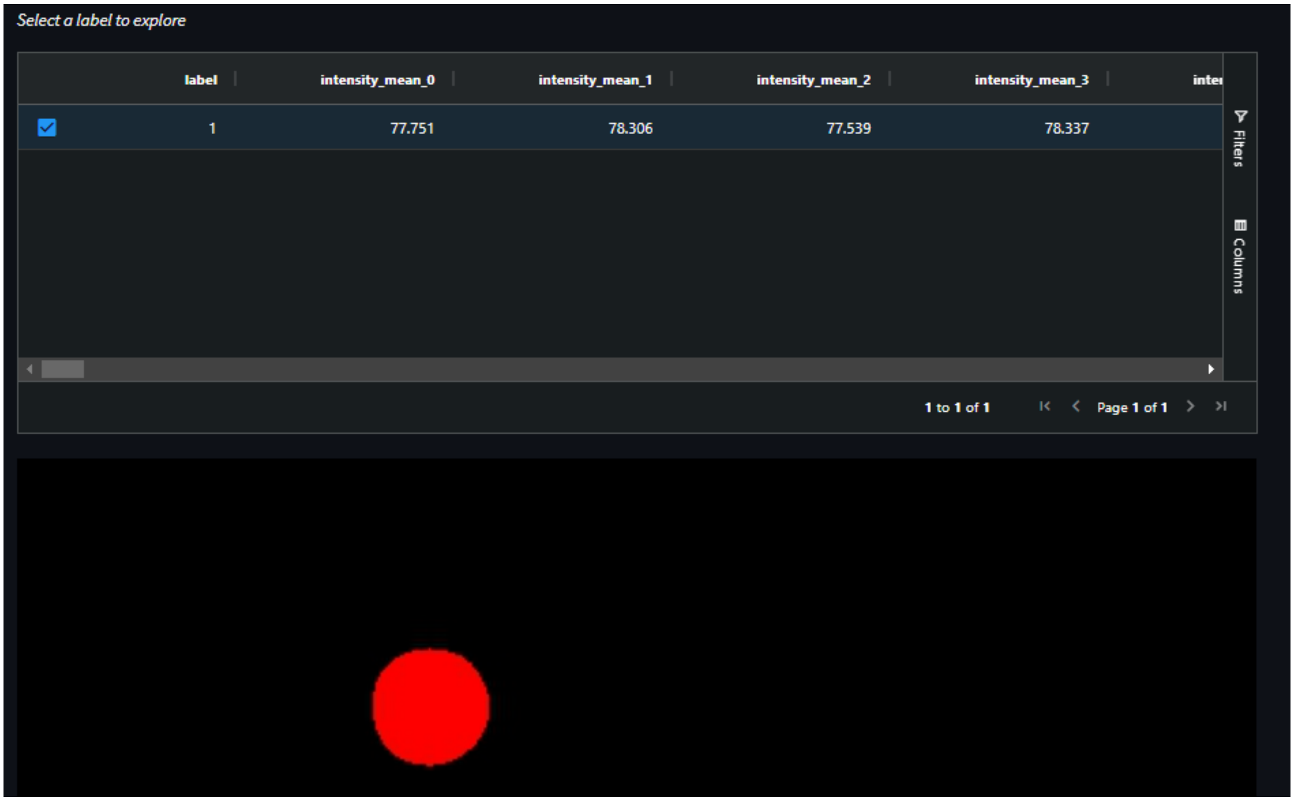

**B**

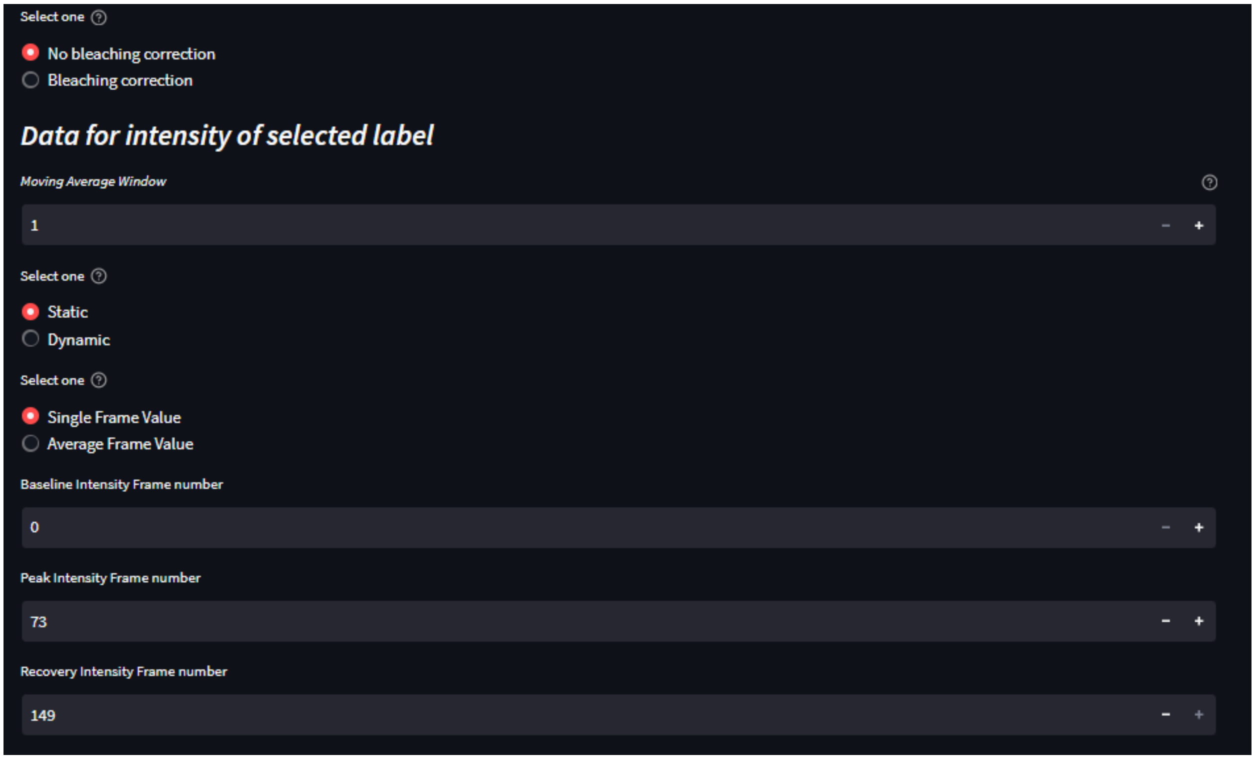
Select one
No bleaching correction
Bleaching correction
Data for intensity of selected label
Moving Average Window
1
Select one
Static
Dynamic
Select one
Single Frame Value
Average Frame Value
Baseline Intensity Frame number
0
Peak Intensity Frame number
73
Recovery Intensity Frame number
149

**C**

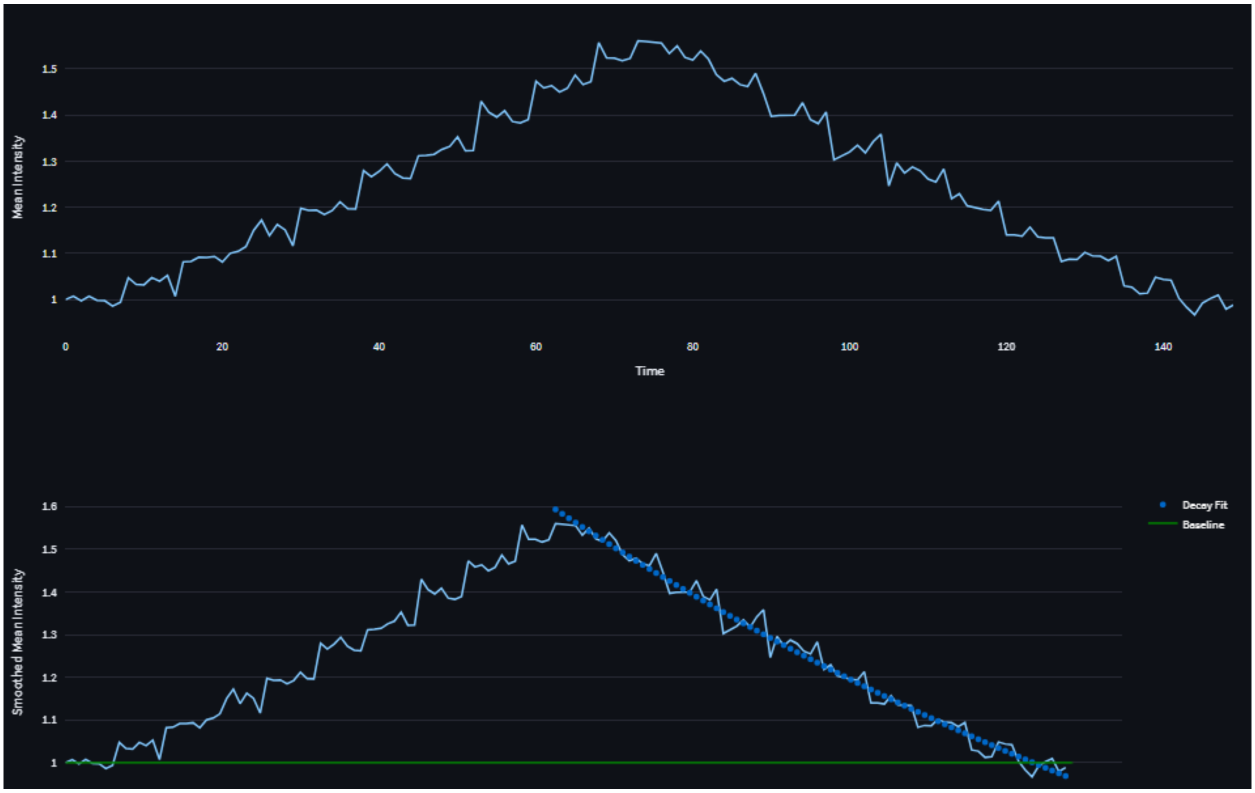
Mean Intensity
1.5
1.4
1.3
1.2
1.1
1
0
20
40
60
80
100
120
140
Time
Smoothed Mean Intensity
1.6
1.5
1.4
1.3
1.2
1.1
1
Decay Fit
Baseline

**Figure S10.** Singe-cell Analysis. (A) A cell selected in an interactive table and highlighted. (B) Options for bleaching correction, trace smoothing, and baseline, peak, and recovery selection. (C) Mean Intensity and Smoothed Mean Intensity traces.

Figure S11 shows a typical example for $Na^+$ ion dynamics in a neuron. With a frame rate of 1, moving average of window 1, and taking average intensity of the first 10 frames for baseline calculation (Dynamic Analysis), the original and the smoothed intensity as functions of time are plotted (Figure S11-A). When the signal is present (the trace rises and decays crossing the baseline), clicking “Obtain the parameters for selected label” computes and outputs various parameters linked to the selected cell (Figure S11-B).

**A**

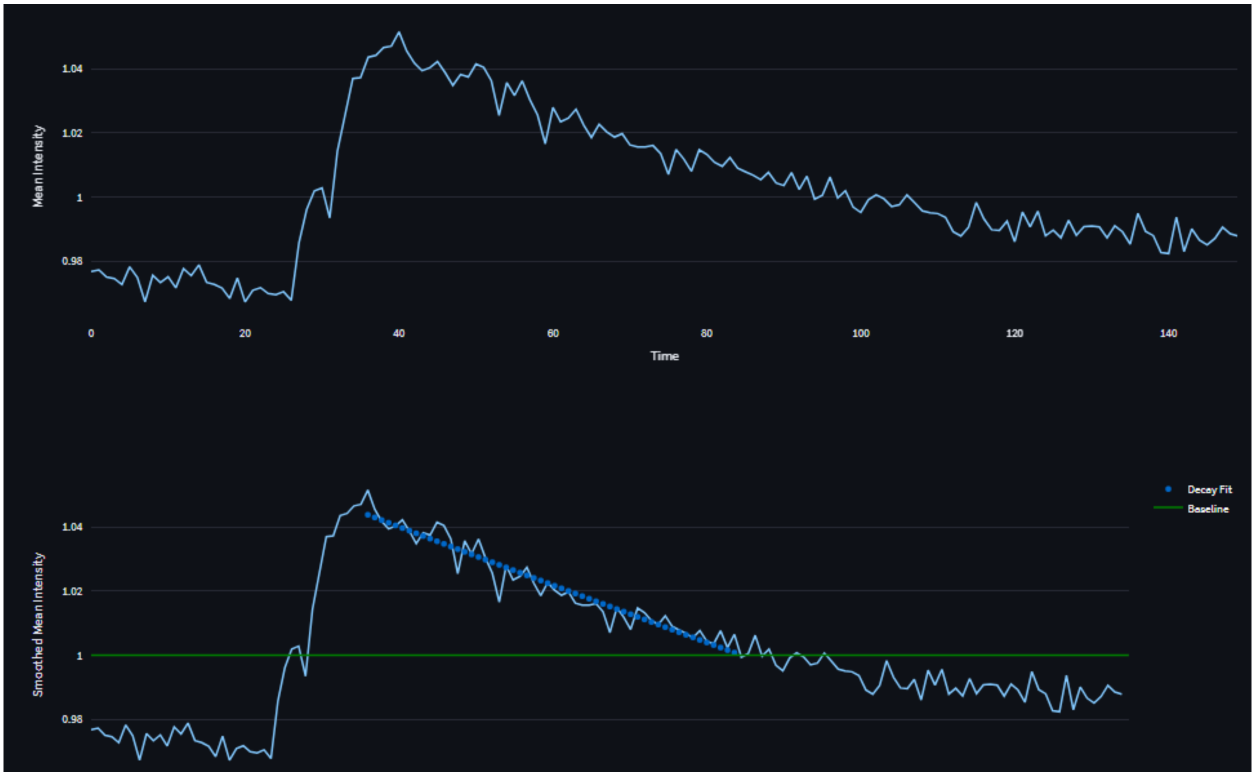

**B**

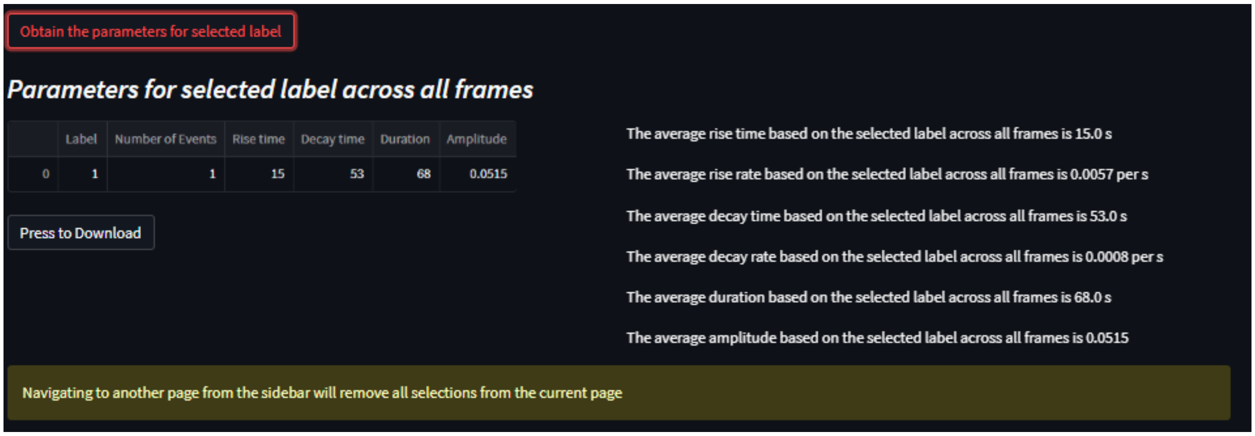

**Figure S11.** (A) Original Mean Intensity and Smoothed mean Intensity (moving average of window 1) as functions of time. The exponential fit to determine the decay rate is shown. (B)

Different properties of the trace computed from the Smoothed mean Intensity curve and displayed.

### Bleaching Correction

When this option is selected, users will get further options to

- Select “Static” or “Dynamic” analysis to be performed (Figure S12).
    1. Adjust the moving average window for trace smoothing, that ranges from 1 to 5 (where 1 would mean the original trace) (Figure S12).
    2. Choose the number of the first and last few frames to fit a mono-exponential curve to correct for photobleaching.

All the other processes and outputs are similar to the “No bleaching correction” option, as shown in Figure S11.

**A**

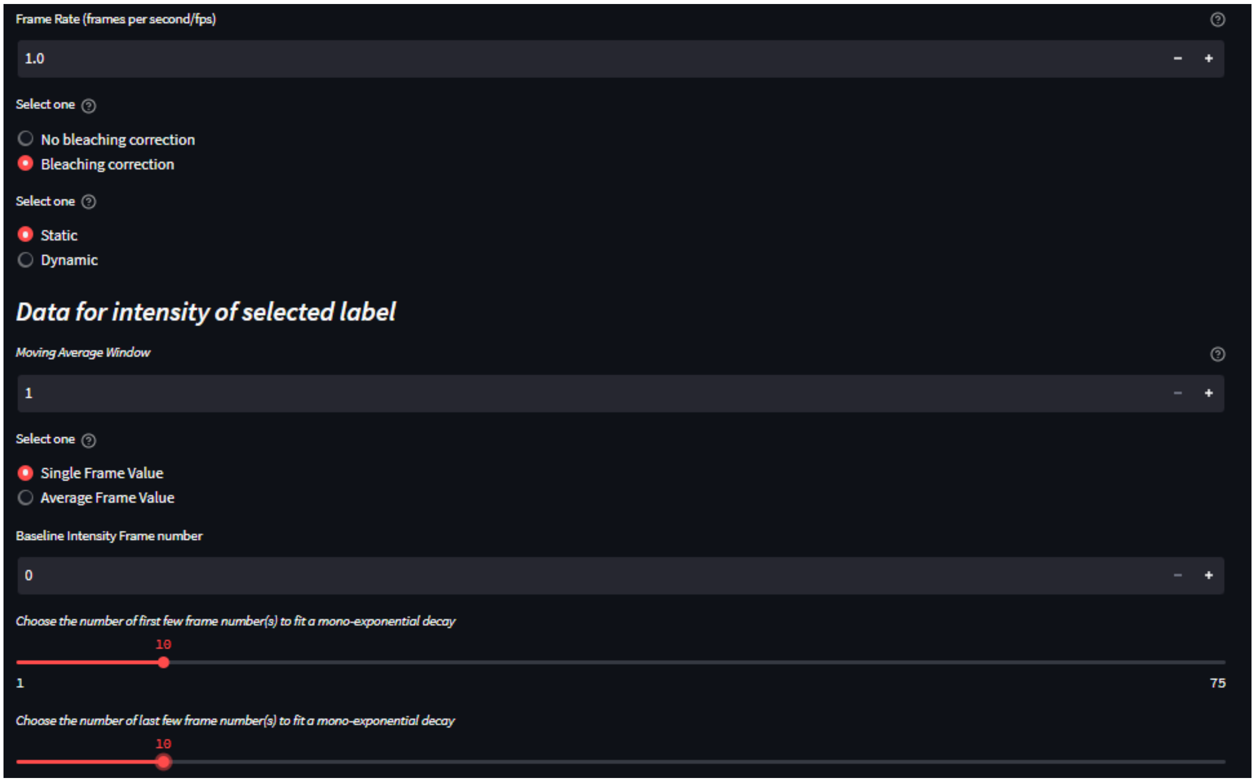

B

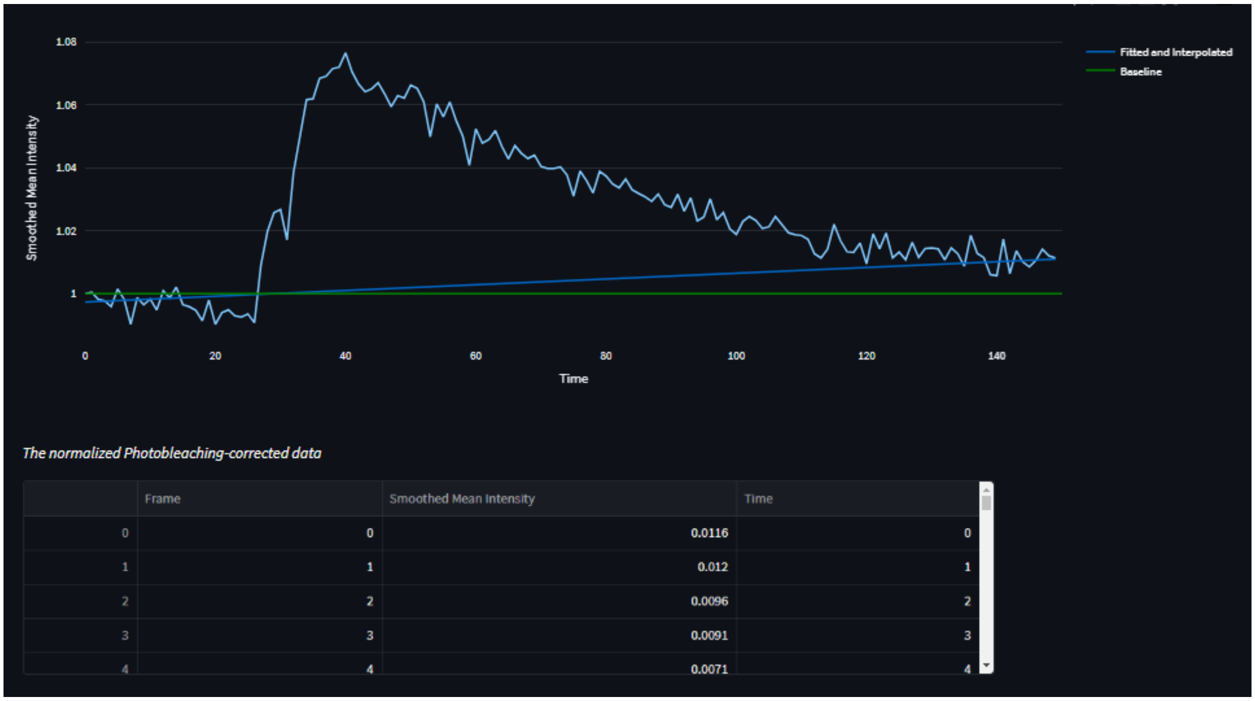

C

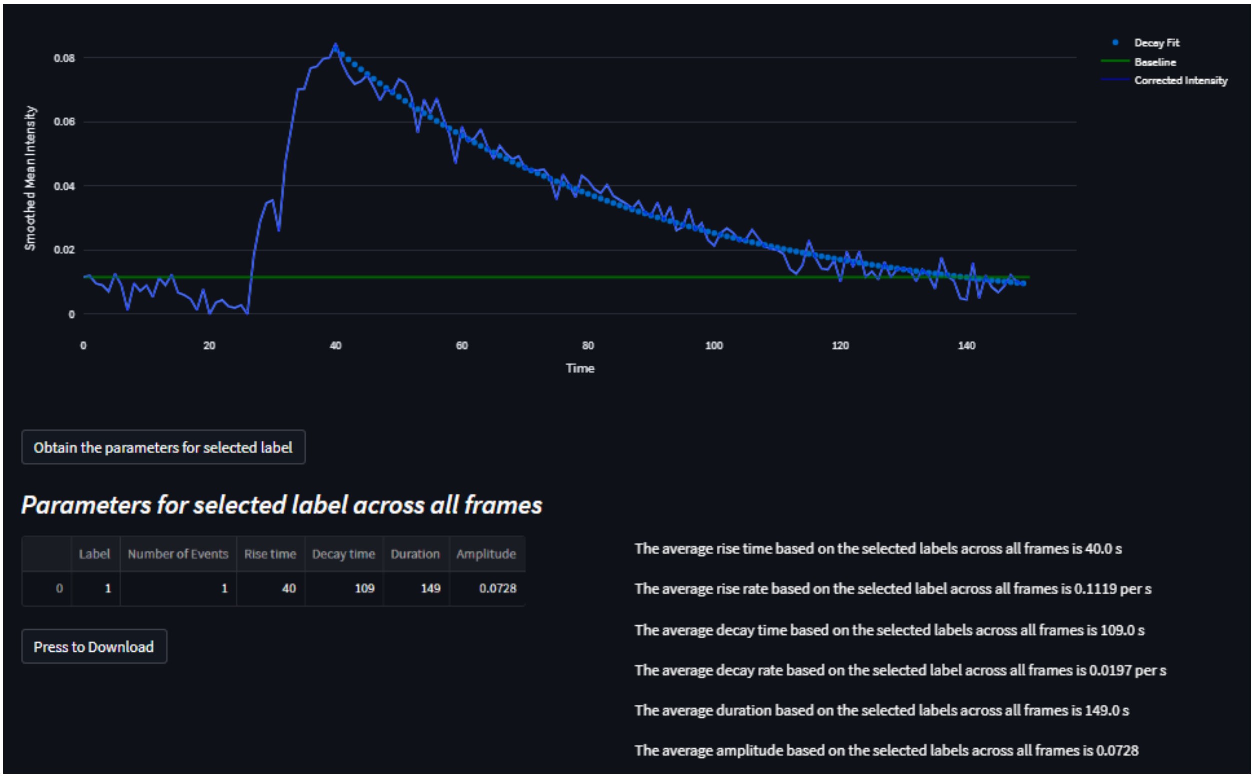

**Figure S12.** (A) Bleaching correction option, followed by the "Static" option selected. (B) The mono-exponential decay curve is fitted using the selected first and last 10 frames, with intermediate points interpolated. The fitted curve is then subtracted from the original intensity to correct for photobleaching. (C) The corrected intensity trace with decay curve for the selected baseline, peak, and recovery frames and the computed parameters.

### F. Multi-cell Analysis

In this section, users can collectively analyze multiple (all or fewer) cells by selecting multiple cells at once. As soon as the cells are selected, their corresponding traces are displayed, which can be isolated by double clicking their legends. All the other options remain unchanged, just as they were for single-cell analysis previously. Clicking “Obtain the parameters for selected labels” displays the normalized traces and computes the parameters based on the selected options (Figure S13).

**A**

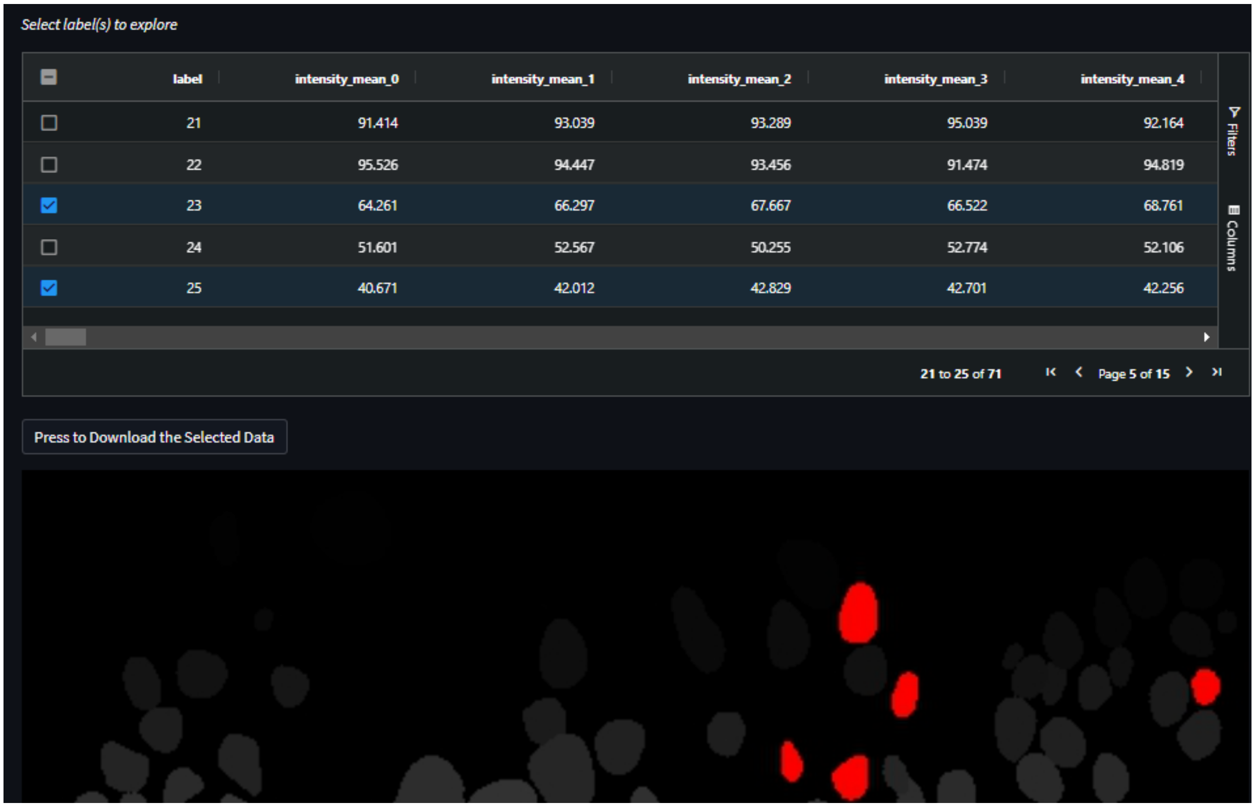

B

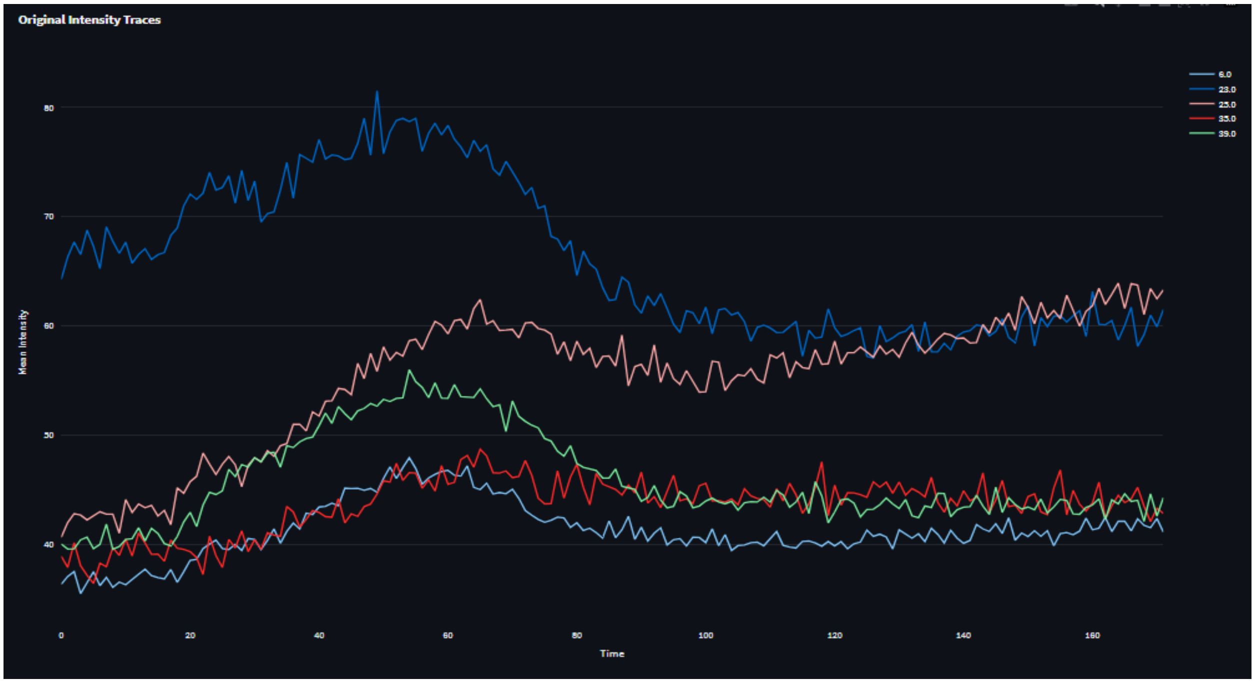
Original Intensity Traces
6.0
23.0
25.0
35.0
39.0
Mean Intensity
80
70
60
50
40
0
20
40
60
80
100
120
140
160
Time

C

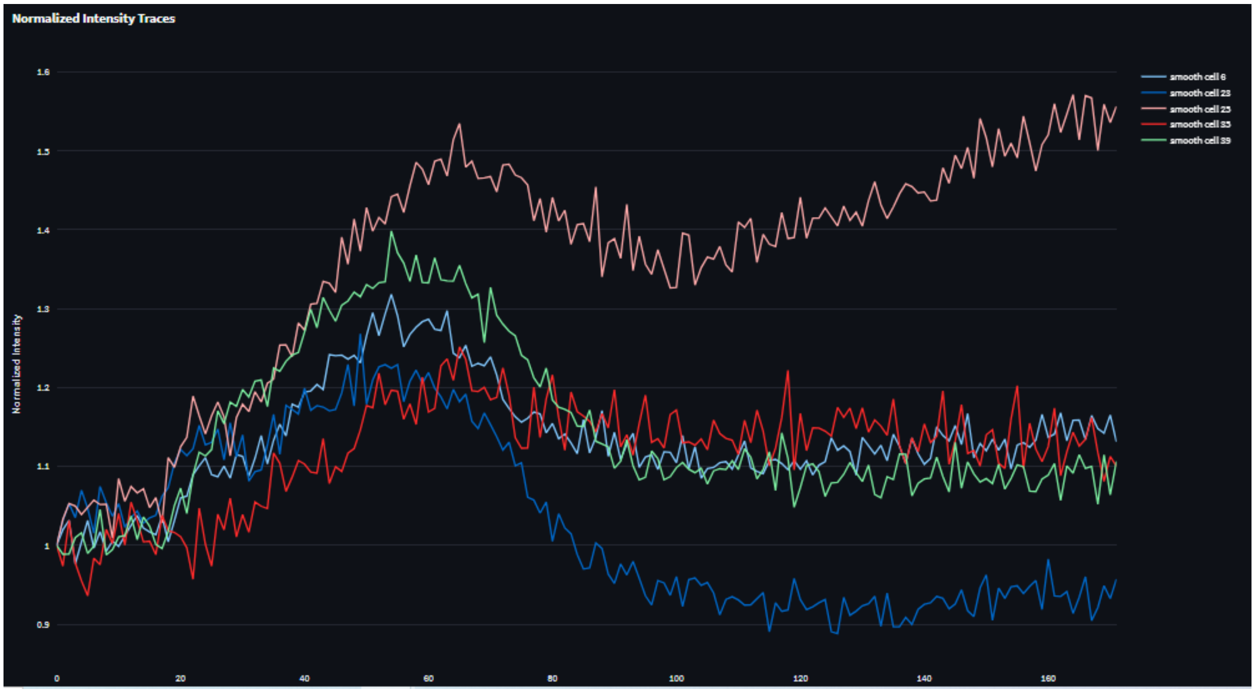
Normalized Intensity Traces
smooth cell 6
smooth cell 23
smooth cell 25
smooth cell 35
smooth cell 39
Normalized Intensity
1.6
1.5
1.4
1.3
1.2
1.1
1
0.9
0
20
40
60
80
100
120
140
160

D

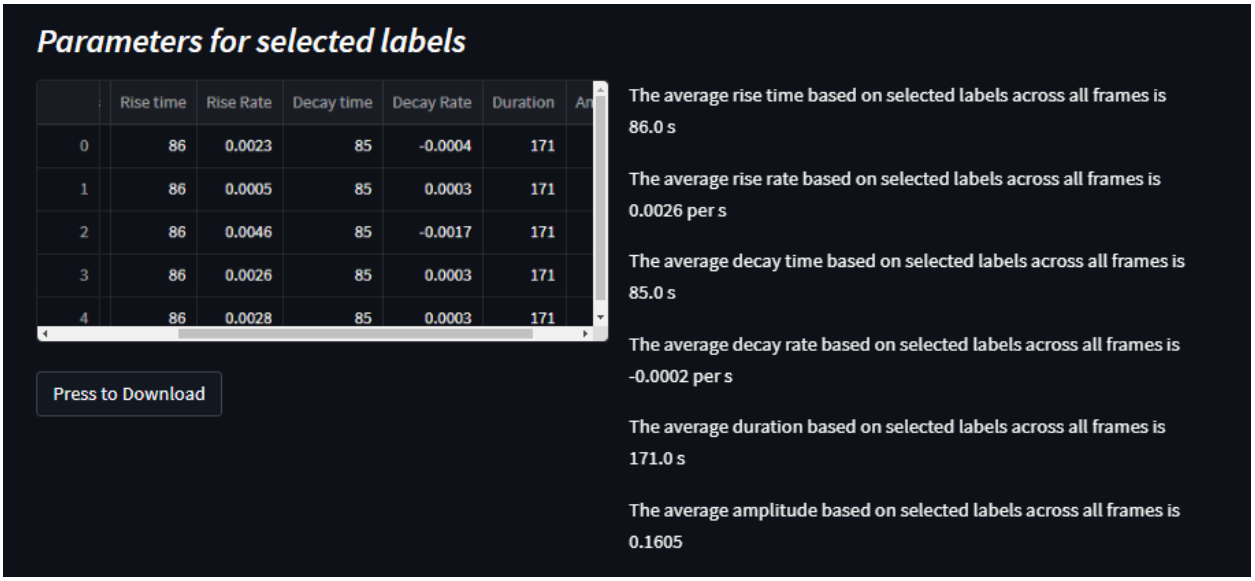

**Figure S13.** Multi-cell Analysis. (A) Multiple cells selected and highlighted on the labeled image. (B) Intensity traces of the selected cells. Double-clicking the label legend isolates the trace. (C) Normalized traces for selected cells. (D) Parameters displayed for selected cells based on the chosen options.

Finally, DL-SCAN generates and displays distribution plots for the computed parameters, as shown in Figure S14.

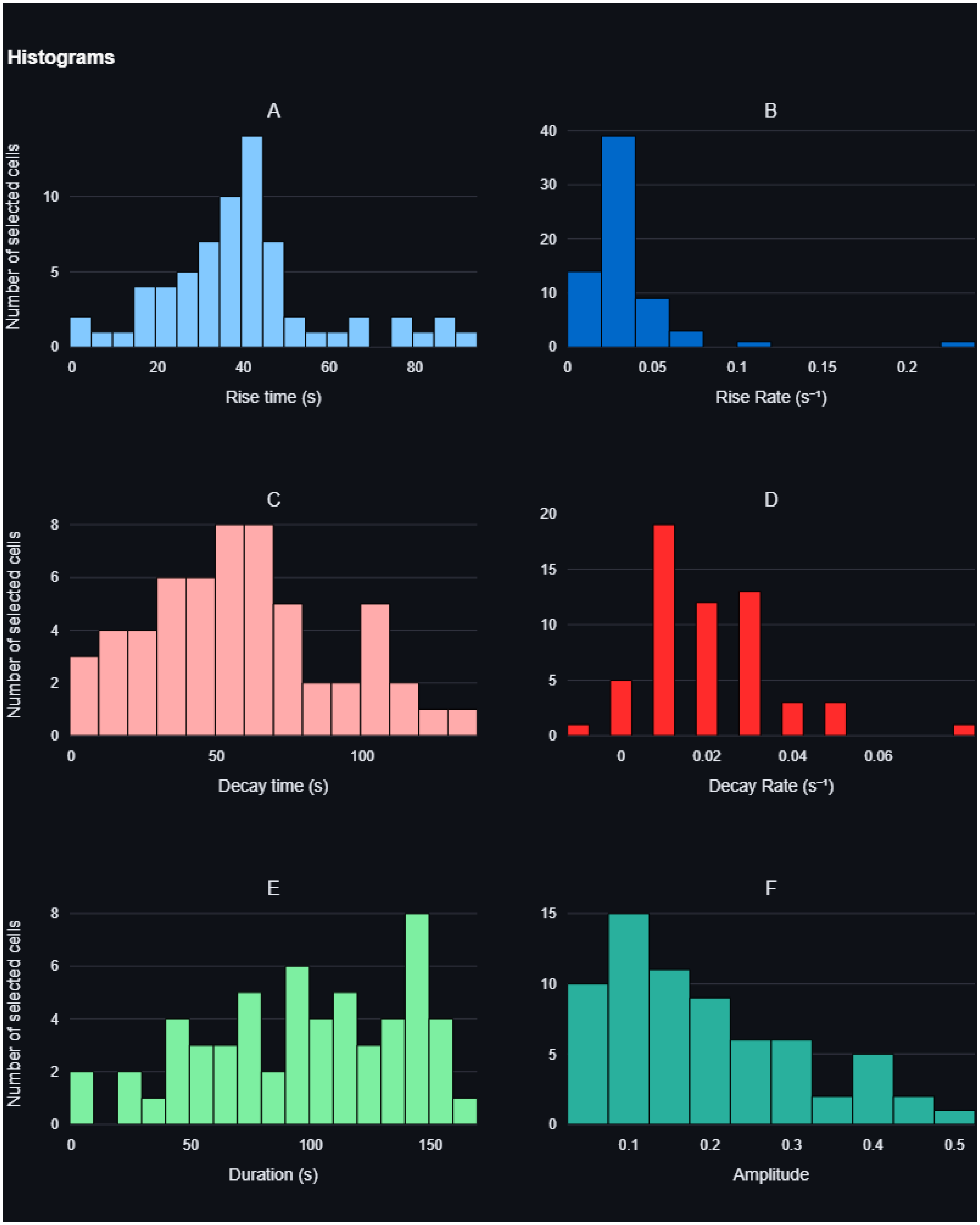

**Figure S14.** Display of distributions of DL-SCAN-generated rise time (A), rise rate (B), decay time (C), decay rate (D), duration (E), and normalized amplitude (F).
